# Rapid Assessment of Biomarkers on Single Extracellular Vesicles Using ‘Catch and Display’ on Ultrathin Nanoporous Silicon Nitride Membranes

**DOI:** 10.1101/2024.04.29.589900

**Authors:** Samuel N. Walker, Kilean Lucas, Marley J. Dewey, Stephen Badylak, George Hussey, Jonathan Flax, James L. McGrath

## Abstract

Extracellular vesicles (EVs) are particles secreted by all cells that carry bioactive cargo and facilitate intercellular communication with roles in normal physiology and disease pathogenesis. EVs have tremendous diagnostic and therapeutic potential and accordingly, the EV field has grown exponentially in recent years. Bulk assays lack the sensitivity to detect rare EV subsets relevant to disease, and while single EV analysis techniques remedy this, they are undermined by complicated detection schemes often coupled with prohibitive instrumentation. To address these issues, we propose a microfluidic technique for EV characterization called ‘**ca**tch and **d**isplay for **l**iquid **b**iopsy (CAD-LB)’. CAD-LB rapidly captures fluorescently labeled EVs in the similarly-sized pores of an ultrathin silicon nitride membrane. Minimally processed sample is introduced *via* pipette injection into a simple microfluidic device which is directly imaged using fluorescence microscopy for a rapid assessment of EV number and biomarker colocalization. In this work, nanoparticles were first used to define the accuracy and dynamic range for counting and colocalization by CAD-LB. Following this, the same assessments were made for purified EVs and for unpurified EVs in plasma. Biomarker detection was validated using CD9 in which Western blot analysis confirmed that CAD-LB faithfully recapitulated differing expression levels among samples. We further verified that CAD-LB captured the known increase in EV-associated ICAM-1 following the cytokine stimulation of endothelial cells. Finally, to demonstrate CAD-LB’s clinical potential, we show that EV biomarkers indicative of immunotherapy responsiveness are successfully detected in the plasma of bladder cancer patients undergoing immune checkpoint blockade.

---

Extracellular vesicles (EVs) are secreted particles of plasma membrane origin that are produced by all cells. The umbrella term ‘EV’ encompasses various classes of particles which can broadly be separated into three groups: exosomes, microvesicles, and apoptotic bodies. Exosomes are ∼50-150 nm in size and are formed *via* inward invagination of late endosomal membranes and released upon multivesicular body fusion with the plasma membrane. Microvesicles are formed as a direct result of membrane budding and are larger in size, ∼150-1000 nm. Apoptotic bodies comprise the largest class (>1 µm) and are released following membrane blebbing during cell death. EVs have gained increased attention over the past few decades, due to a number of reasons. Foremost, EVs carry bioactive cargo, including proteins, nucleic acid, and other soluble factors from their parent cells that facilitate intercellular communication.^1–3^ EVs are found at high levels in biofluids including blood, urine, saliva, cerebrospinal fluid, breast milk, amniotic fluid, semen, and bile.^4^ Thus, EVs exhibit diagnostic value since they represent circulating biomarkers that can be evaluated as a liquid biopsy tool. Furthermore, EVs carry out defined functional roles in normal physiology and disease pathogenesis.^3, 5–7^ For example, EVs can drive cancer progression by aiding in metastasis,^8^ conferring chemoresistance,^9^ and modulating immune responses.^10^ Accordingly, EVs have also been studied as a therapeutic target with the goal of controlling native EV trafficking or using biomimetics for therapeutic agent delivery.^11, 12^

Despite the tremendous potential of EVs, their analysis is nontrivial. EVs are vastly heterogeneous and various unique subsets can be identified in the secretome of single cells.^13, 14^ Furthermore, the EV field is still relatively new and standardized procedures for EV isolation and characterization do not exist. EV isolation and subpopulation sorting are accomplished using differential centrifugation, size-based, immunoaffinity capture, polymer precipitation, and/or microfluidic techniques.^15^ Enrichment methods add considerable time to EV analysis and inevitably result in the loss of unique EV subsets. Nanoparticle tracking analysis (NTA) is most commonly used to determine EV concentration, although there is evidence that reported concentrations are over-estimations and highly dependent on user settings.^16–20^ Moreover, NTA instruments offer a limited dynamic range; sample concentration is essentially confined within a single order of magnitude.^18^ EV cargo has traditionally been studied using Western blot or ELISA analyses. These bulk assays pool large numbers of EVs and provide biomarker information averaged across the pooled sample. In liquid biopsy assays, EV signals from target cells are thought to be extraordinarily low relative to the high background of EVs secreted from other cells throughout the body. Bulk assays lack the sensitivity required to detect rare EV subsets, and thus, single EV technology is necessary for diagnostic applications. Numerous single EV analysis methods exist including nano-flow cytometry (nFC),^21^ microscopy,^22^ digital PCR,^23^ digital ELISA,^24^ Raman spectroscopy,^25^ plasmonics,^26^ NTA,^27^ and others.^28^ Some single EV techniques demonstrate remarkable sensitivity but rely on complicated detection schemes that offer limited throughput,^22–24, 26^ or suffer from the inability to interrogate multiple biomarkers simultaneously.^25, 27^ Biomarker multiplexing is critical to determine cell-of-origin identity and functional state.

nFC represents the leading candidate for a standardized EV analysis approach,^21, 29^ however, instrumentation is expensive, often requires careful development, and operators must develop sophisticated expertise.^30^ A large gap exists for single EV technology, namely, current techniques don’t translate to clinical diagnostic settings. For this, methods must demonstrate rapid throughput and reproducibility, and enable wide adoption through a user-friendly manufacturable format.^31^ Accordingly, here we introduce an EV assay that operates in a simple microfluidic device that can be rapidly assembled from mass-produced components. This device features an ultrathin (∼100 nm) nanoporous silicon nitride (NPN) membrane whose pores are appropriately sized and spaced to capture individual EVs. Because NPN membranes are ultrathin and inorganic, they are extraordinarily permeable,^32, 33^ compared to conventional polymeric membranes, and optically transparent by light microscopy.^34^ The feasibility of NPN membrane EV capture has been demonstrated previously.^35, 36^ NPN membranes have also been used to study single EV functionality *via* maintenance of luminal pH by the sodium-hydrogen exchanger NHE1.^37^ The µSiM (**micro**fluidic device featuring an ultrathin **si**licon **m**embrane) is a device platform originally developed for cell culture applications.^38^ The µSiM platform has also been used as a filter-based capture tool for diagnostic applications in an adaptation we term the ‘µSiM-DX’.^39^ Here the µSiM- DX strategy is reconfigured to generate a pipette-powered microfluidic scheme for single EV analysis we call ‘**ca**tch and **d**isplay for **l**iquid **b**iopsy (CAD-LB)’.

As an imaged-based tool that enables isolated EV and biomarker enumeration, CAD-LB is a ‘digital’ assay. While they typically require complex workflows, digital assays have advantages in both sensitivity and dynamic range compared to bulk assays. Specifically, single-analyte imaging-based immunoassays have enhanced signal detection by digitally counting target analytes.^40–42^ A common approach is forming analyte-microbead conjugates which facilitate target analyte capture for direct imaging^40^ or ELISA detection.^41^ The development of single EV imaging platforms has paralleled that of conventional protein assays, but methods require both extensive microfluidic manipulation and reagent modification to isolate individual EVs and generate capture immunocomplexes, respectively.^23, 24^ Direct imaging of surface-captured EVs typically requires surface modification to both EVs and the capture surface.^22, 43^ Imaging single EVs in solution is possible but this requires tracking algorithms and EV diffusion can obscure multiplexed biomarker detection.^44^ Common to all digital EV techniques is considerable upstream processing; reliable signal detection requires time-consuming EV isolations and post-labeling washes or purification.^22–24, 43, 44^ Unlike all aforementioned methods, CAD-LB rapidly isolates hundreds of thousands of single EVs from minimally processed samples and allows the passage of contaminating protein and antibody molecules. We demonstrate here that NPN membranes are an ideally structured surface that isolates individual EVs into pores spaced at the limits of conventional microscopy resolution (∼100 nm). While we focus here on assessing CAD-LB accuracy for counting EVs and colocalized biomarkers using confocal microscopy, our report provides the foundation for future work that employs super-resolution techniques to survey hundreds of millions of single analytes to discover instances of very rare biomarkers.

## RESULTS & DISCUSSION

### ‘Catch and Display for Liquid Biopsy (CAD-LB)’ Platform

A schematic depicting the general CAD-LB workflow is presented in **Figure 1**. First, solutions containing EVs are fluorescently labeled using a pan EV label (carboxyfluorescein succinimidyl ester (CFSE)) and fluorescently-conjugated antibodies targeting EV surface proteins of interest (**Figure 1: Left Panel**). The key material is an ultrathin NPN membrane with roughly 100 million ∼60 nm pores (∼15% porosity).^32^ At this pore size, NPN captures EVs while allowing for the passage of monomeric proteins at concentrations as high as 10 mg/mL.^45^ Thus, unreacted antibody molecules and protein contaminants are cleared during capture (**Figure 1: Middle Panel**) and do not need to be removed in a separate step. The ultrathin nature of NPN membranes makes them extraordinarily permeable compared to conventional polymeric membranes. This enables microfluidic filtration at low pressures^35^ and produces a dense field of captured EVs if the membrane is used to full capacity (pores on NPN are spaced ∼131 nm apart). The membrane thinness provides optical transparency and their inorganic Silicon Nitride composition gives no autofluorescence at visible wavelengths. These qualities make NPN a glass-like, nanostructured material that is ideal for capturing isolated EVs for imaging by fluorescence microscopy (**Figure 1: Right Panel**). The resulting images can then be analyzed for both EV quantification and colocalization with protein biomarkers.

**Figure 1.**
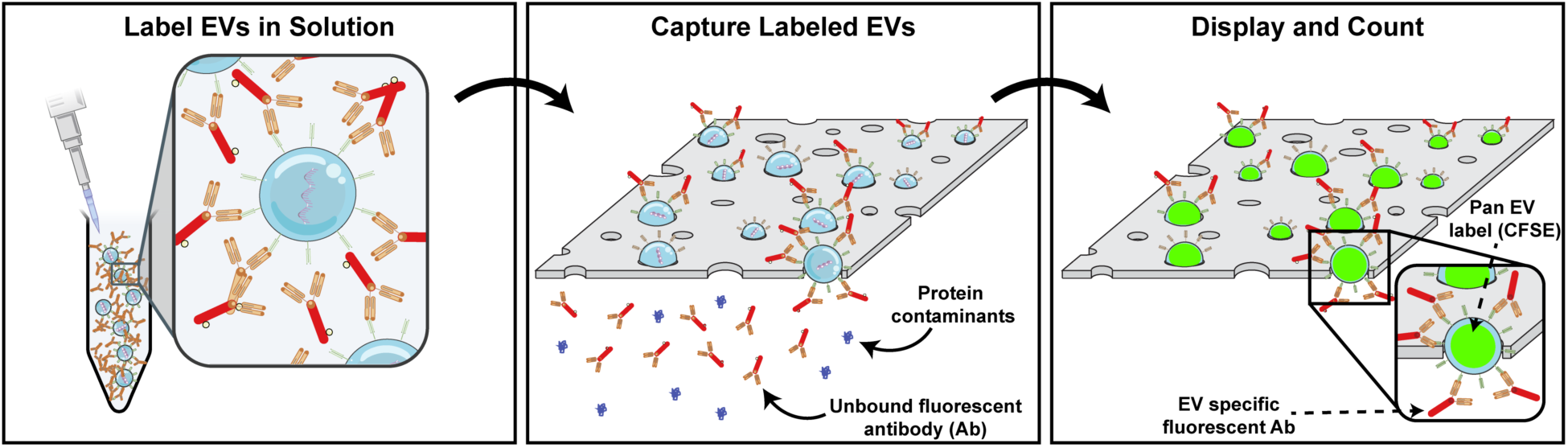
The ‘Catch and Display for Liquid Biopsy (CAD-LB)’ Platform. Schematic of single extracellular vesicle (EV) capture and detection on ultrathin nanoporous silicon nitride (NPN) membrane. (Left) EVs are fluorescently labeled in solution with carboxyfluorescein succinimidyl ester (CFSE) and an antibody targeting a biomarker of interest **(Middle)** This solution is then passed through a simple microfluidic device where EVs are captured within membrane pores while unreacted antibody molecules and protein contaminants are filtered through the NPN membrane. **(Right)** The membrane is then imaged using fluorescence microscopy to identify CFSE signals and antibody signal colocalization indicating biomarker-positive EVs.

### Microfluidic Device Development

To enable CAD-LB we sought a microfluidic workflow that achieves EV capture in one step and enables direct fluorescence imaging in the same device without any further processing. To start, we explored the use of the **t**angential **f**low for **a**nalyte **c**apture (TFAC) filtration device developed previously by our team for nanoparticle and EV imaging by scanning electron microscopy (SEM) (**Figure 2A**).^35, 36^ The TFAC device features microfluidic channels above and below the NPN membrane and is hand-assembled in a complex and labor-intensive, layer-by-layer fashion. The two microfluidic channels allow for precise and independent control over the rate of sample supply (Q_s)_ and ultrafiltration through the membrane (Qu) (**Figure 2B**). Under this flow regime, species slightly larger than NPN pores are physically captured during ultrafiltration while small contaminants pass through. Very large species rejected by the membrane are cleared from the membrane surface by the tangential flow component, thus ameliorating fouling.^46^ In our prior work, NPN membranes were retrieved from disassembled devices for imaging by SEM. SEM micrographs showed single EV capture per pore and nanogold-conjugated anti-CD63 antibody labeling on single EVs from undiluted plasma samples.^35^

**Figure 2.**
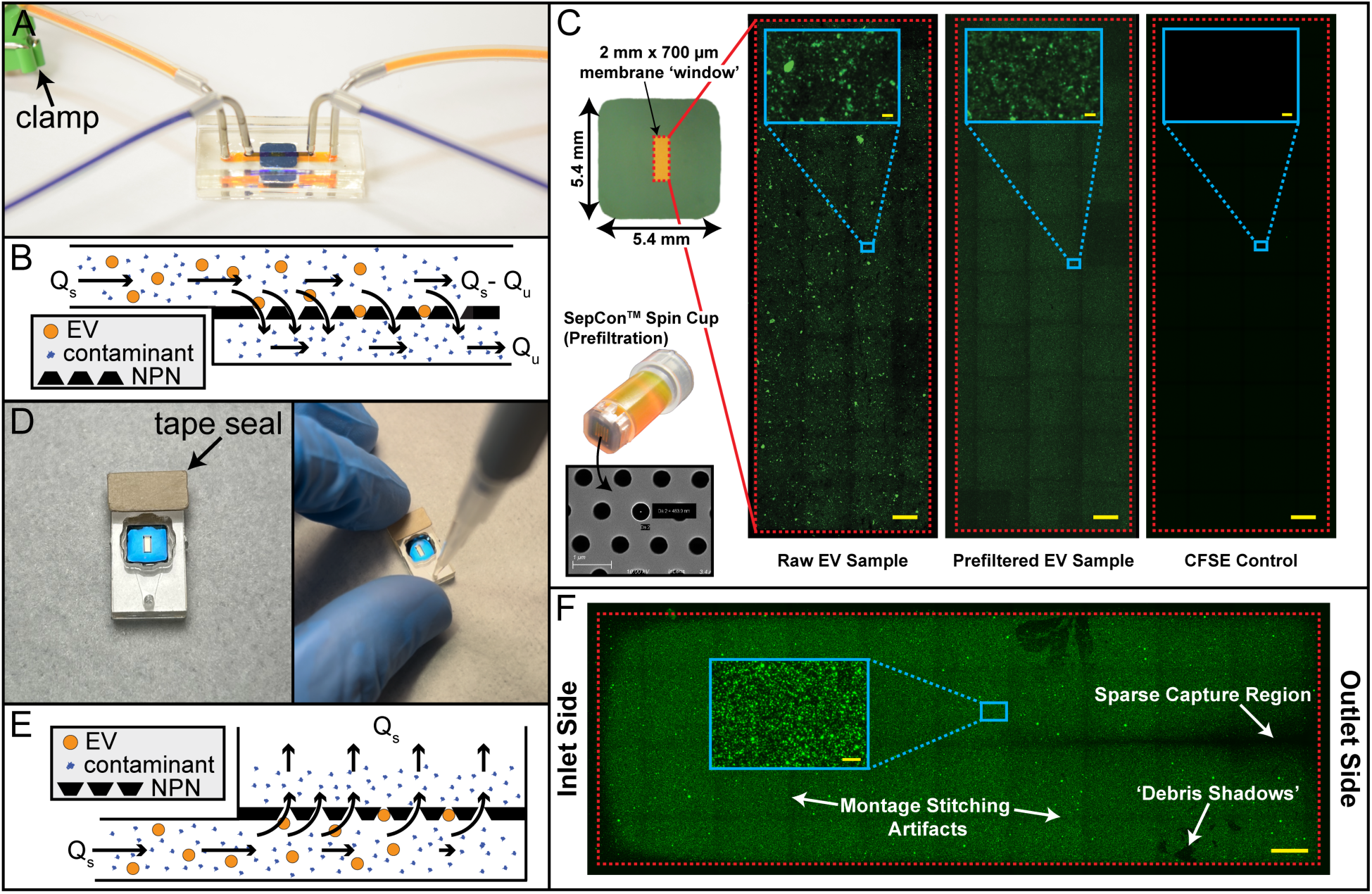
CAD-LB Device Development. **(A)** First generation CAD-LB hand-built device fashioned using layer-by-layer assembly and operated under tangential flow for analyte capture (TFAC) flow regime. **(B)** Cross-section device schematic depicting TFAC flow regime. **Qs** and **Qu** represent supply and ultrafiltration flow rates, respectively, maintained by syringe pumps. **(C)** Montage fluorescence microscopy images of carboxyfluorescein succinimidyl ester (CFSE)-labeled extracellular vesicles (EVs) collected from urinary bladder cancer grade II (5637) cell line captured on ultrathin nanoporous silicon nitride (NPN) membranes using TFAC devices. Montage images include unfiltered raw EVs **(left),** EVs prefiltered with SepCon™ spin cups **(middle),** and CFSE-only control **(right).** Scale bars = 100 pm, inset scale bars = 5 pm. **Bottom Left Inset:** SepCon™ spin cup used for prefiltration, spin cups contain microporous membranes with 0.5 pm pores. **(D)** Second generation CAD-LB device incorporates modular pSiM (microfluidic device featuring an ultrathin silicon membrane) platform. Blocking one microfluidic channel port **(left)** creates a dead-end-tangential compound flow regime sustained *via* pipette injection **(right). (E)** Cross-section device schematic depicting dead-end-tangential compound flow regime. **Qs** denotes supply flow rate sustained by pipette injection. (F) Montage fluorescence microscopy image of 100 nm fluorescent nanoparticles captured on NPN membrane using modified pSiM device. Scale bar = 100 pm, inset scale bar = 5 pm.

To extend the TFAC method to allow for fluorescence detection of EVs by optical microscopy, EVs isolated from conditioned cell culture (5637) media were labeled with the luminal EV marker CFSE. Devices again needed to be disassembled and membranes retrieved to enable high- resolution imaging using a 60X objective because TFAC devices would otherwise require a working distance >1 mm for *in situ* imaging. Membranes transferred to glass bottom dishes were imaged in their entirety on a spinning disk confocal microscope by montaging 40 fields of view (FOVs) (**Figure 2C**). Despite the need for device disassembly and membrane manipulation prior to imaging, the dense capture field indicates uniform EV capture and very high retention rates which is supported by our previous findings where captured gold nanoparticles were imaged by SEM.^36^ To reduce the polydispersity of CFSE-positive material in conditioned media samples, we added a short prefiltration step using a microcentrifuge and SepCon^TM^ spin cups (0.5 µm membrane filter) (**Figure 2C: Bottom Left Inset**). Prefiltration cleared larger aggregates/debris and produced a uniform field of smaller CFSE-labeled particles.

As a second option for the microfluidic device, we examined the µSiM platform recently developed by our laboratory for cell culture applications.^38^ Unlike the TFAC device, the µSiM is rapidly assembled from mass-produced components lined with pressure-sensitive adhesives that are activated during assembly in alignment jigs. The µSiM features a lower microfluidic channel and an upper open well separated by an ultrathin silicon membrane. Klaczko *et al.* used the µSiM for viral detection by injecting 40 µL of sample into the microfluidic channel using only a pipette.^39^ The membranes used in these devices were functionalized with a capture protein and featured micron-sized pores. Upon injection the sample followed the path of least hydraulic resistance, filtering through the membrane and into the well. When intact virions were present, their affinity-based capture resulted in membrane ‘clogging’ and a switch in the path of least resistance; the sample rerouted through the microfluidic channel and exited the outlet port. The use of the µSiM platform for diagnostics *via* filtration and species capture was then termed the ‘µSiM-DX’.

To employ the same pipette-powered injection strategy as Klaczko *et al.*, but with a nanoporous membrane, we first needed to block the outlet port of the microfluidic channel with strong adhesive tape (**Figure 2D**). Otherwise, the much higher hydraulic resistance of the NPN membrane would simply route the injected sample under the membrane to the outlet port. Importantly, because the working distance for the µSiM-DX is only ∼300 µm, it is possible to directly image the membrane with a 60X objective and avoid the need for device disassembly and membrane manipulation to enable microscopy. Because the flow pattern in the µSiM-DX with a blocked port is a complex mix of dead-end and tangential flow filtration, which changes from the inlet to the outlet side of the membrane (**Figure 2E)**, the resulting capture field exhibits features distinct from TFAC (**Figure 2F**). Specifically, the flow bifurcates and creates a small region of sparse capture (<1% of total membrane area) near the membrane midline on the outlet port side. Also, since the tangential flow component diminishes further from the injection port, large debris can be captured on the membrane which appears as dark shadows in the capture field (**Figure 2F**). While there are benefits to TFAC for capture profile clarity, the facile device assembly, easy operation, low sample volume requirements, and *in situ* imaging benefits of the µSiM-DX make it our preferred device for CAD-LB.

### Evaluating the Accuracy & Dynamic Range of CAD-LB

Having selected a device platform, we evaluated the quantitative performance of CAD-LB with 100 nm fluorescent nanoparticles serving as surrogates for EVs. Nanoparticle inputs spanning six orders of magnitude (10^2^ - 10^7^) were prepared according to the manufacturer specifications and processed with the CAD-LB workflow in triplicate. This range of concentrations was selected to end near full capacity of the membrane (*i.e.* one nanoparticle for each pore) and starts at just ∼0.0004% of the total capacity. Confocal microscope images of the membrane were analyzed with FIJI to count the number of captured nanoparticles (see Experimental Section). Counts were then compared to expectations based on the injected number of nanoparticles with a correction for an observed systematic error likely attributed to loss upstream of capture (**Figure S1**).

**Figure 3** illustrates the experimental results and accompanying error analysis for the nanoparticle studies. Images can be appreciated to transition from those featuring clearly isolated nanoparticles at an input of 7⋅10^4^ to a highly crowded field at an input of 7⋅10^6^ (**Figure 3A-C**). CAD-LB exhibits high reproducibility at every input and a highly linear response between 10^2^ -10^6^ (R^2^ = 0.976; **Figure 3D**). Beyond 10^6^, increasing nanoparticle crowding caused a progressively worse undercount. Expressing error in terms of log_10_ demonstrates that CAD-LB faithfully reports the order of magnitude for all inputs within the linear range (**Figure 3E**). For inputs greater than 10^6^, the error rises due to undercounting and eventually reaches a full decade as the loading approaches the membrane’s capacity. Considering the error in terms of average interparticle distance at each input, the largest error occurs when nanoparticles surpass the optical resolution of our imaging system (∼306 nm) and approach the size of camera pixels (108 nm) (**Figure 3F**). Clearly, super-resolution imaging should be able to extend the dynamic range, however, here we assess a CAD-LB workflow that only requires basic instrumentation and enables wide use. For the current imaging system, our analysis suggests that we limit particle density to 10^6^ or less. It is worth noting that undercounting error begins at lower inputs than we would expect if the only contribution were the limits of the optical system. We attribute the onset of undercounting to a different phenomenon where, due to a non-uniform distribution of pore sizes, shapes, and spacing, adjacent nanoparticles can be captured within the optical limits even at lower inputs. If both the NPN pore dimensions and spacing were more uniform, the error seen for inputs in the 10^6^ range would be reduced to extend the linear quantification region by another decade.

**Figure 3.**
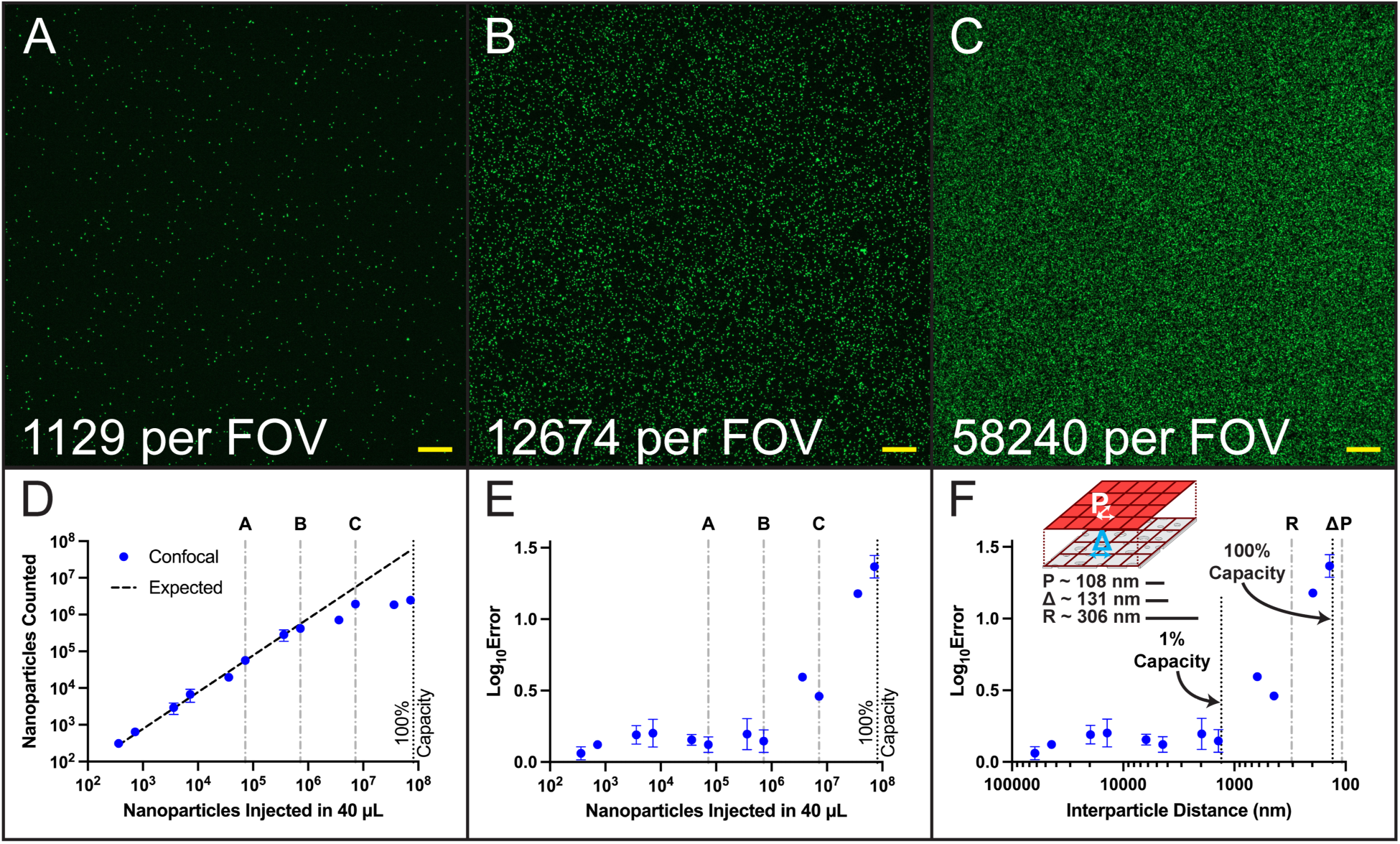
Nanoparticle Capture Experimental Results & Error Analysis. **(A-C)** Representative fields of view (FOVs = 206 pm x 206 pm) from confocal microscope images taken following 100 nm fluorescent nanoparticle capture on ultrathin nanoporous silicon nitride (NPN) membranes. Nanoparticle counts were determined using an open-source FIJI plugin (ComDet v.0.5.5) and are given for respective FOVs in bottom left corner of panels. **(A), (B),** and **(C)** correspond with theoretical nanoparticle inputs of 7▪ 10^4^, 710^5^, and 7-10^6^, respectively. Scale bars = 15 pm. **(D)** Comparison of nanoparticles counted and theoretical nanoparticle input injected in CAD-LB devices. Total nanoparticle count is extrapolated from the average of 3 FOVs and based on total membrane area. The expected count is adjusted based on a systematic error observed in nanoparticle capture experiments **(Figure S1).** The labels **A-C** denote nanoparticle inputs associated with images shown in panels (A-C). 100% capacity vertical line indicates the case in which the nanoparticle input would be equal to the total number of NPN pores. (E) Counting error presented in terms of |log_10_(expected nanoparticle count) - log_10_(captured nanoparticle count)|. (F) Comparison of counting error and theoretical interparticle distance following capture. The dimensions of the optical resolution of the imaging system **(R = 306 nm),** average NPN interpore spacing **(A = 131 nm),** and camera pixel size **(P = 108 nm)** are denoted with vertical lines. 1% and 100% capacity vertical lines indicate interparticle distance at respective nanoparticle inputs. **Top Inset:** Cartoon schematic of NPN membrane with approximate camera pixel array overlaid. Relative dimensions of pixel size (P), average interpore spacing (A), and optical resolution (R) are shown with respect to schematic. All captured nanoparticles were imaged with a fluorescence microscope and quantified using an open-source FIJI plugin (ComDet v.0.5.5). All data points represent mean ± SEM (n = 3).

It is instructive to compare the quantification capabilities of CAD-LB to existing tools for EV quantification. In nanoparticle experiments, we emphasize that CAD-LB possesses superior analytical sensitivity (∼10,000-fold greater) and a wider dynamic range (∼69-fold greater) compared to NTA^18^ (**Figure S2**), the standard EV quantification tool. Compared to other commonly used EV quantification instruments, tRPS and nFC, CAD-LB exhibits a nanoparticle dynamic range roughly 35-fold and 140-fold greater, respectively.^18^ Additionally, CAD-LB’s nanoparticle dynamic range is at least an order of magnitude greater than recently developed single EV analysis techniques.^21, 44, 47^ These features highlight the utility of CAD-LB, namely, the ability to work with a broader range of samples and the ability to detect rarer subsets.

### Nanoparticle Colocalization Control Experiments

We next sought to define the limits of CAD- LB to detect colocalized CFSE-labeled EVs and antibody-labeled biomarkers (**Figure 1: Right Panel**). In these experiments, the total antibody signal will be comprised of specific EV labeling events (true positives) as well as disassociated antibody complexes large enough to be captured by NPN pores (antibody molecules pass freely through the pores of NPN).^48^ Just as counting becomes inaccurate at high capture densities, a false colocalization of pan EV and unassociated antibody signals will be increasingly likely at high concentrations. To develop guidelines to help avoid misinterpreting false colocalization, we performed CAD-LB with two nanoparticle species, each labeled with a distinct fluorescent dye (red or blue) and exclusively visualized in separate fluorescence microscope channels. These species representing fully independent EV and antibody markers were mixed in a controlled manner to elucidate rates of false positive colocalization detection as a function of red and blue nanoparticle input.

In these experiments, the red nanoparticle input (pan EV label surrogate) was held constant (10^5^), while the blue nanoparticle input (disassociated antibody label surrogate) spanned four orders of magnitude (10^2^ - 10^5^) (**Figure 4A-F**). Relative to the 10^5^ red input, the rate of false positive colocalization scaled linearly with the blue input (10^2^ : 0.006%, 10^3^ : 0.050%, 10^4^ : 0.482%, 10^5^ : 4.759%) (**Figure 4G**). This finding is expected and enumerates the potential for false positive colocalization in CAD-LB. As a check, we also calculated the false positive rate relative to the blue input and found that it was very consistent: ∼4% (**Figure 4G**). This finding is also expected since the probability of capturing a blue nanoparticle close enough to a red nanoparticle to induce colocalization should depend only on the degree of pore occupancy by red nanoparticles which was held constant. The false positive rates relative to the red nanoparticle background are also presented in terms of blue:red nanoparticle input ratio (**Figure 4H**). In addition to analyzing false positive rates as a measure of CAD-LB specificity, we performed experiments to confirm our ability to detect true positives when signals originate from the same nanoparticle. Here, TetraSpeck^TM^ microspheres, which are labeled by the manufacturer with multiple fluorescent dyes, were prepared at a single input (10^5^) and processed using CAD-LB (**Figure 4I-K**). The true positive detection rate, or colocalization sensitivity, was ∼98.3% suggesting a false negative rate of ∼1.7% (**Figure 4H**). Other single EV technologies have evaluated their intrinsic specificity and sensitivity using nanoparticle mixtures. Compared to these, CAD-LB exhibits superior colocalization specificity with comparable sensitivity^43^ or superior colocalization sensitivity with comparable specificity.^44^

**Figure 4.**
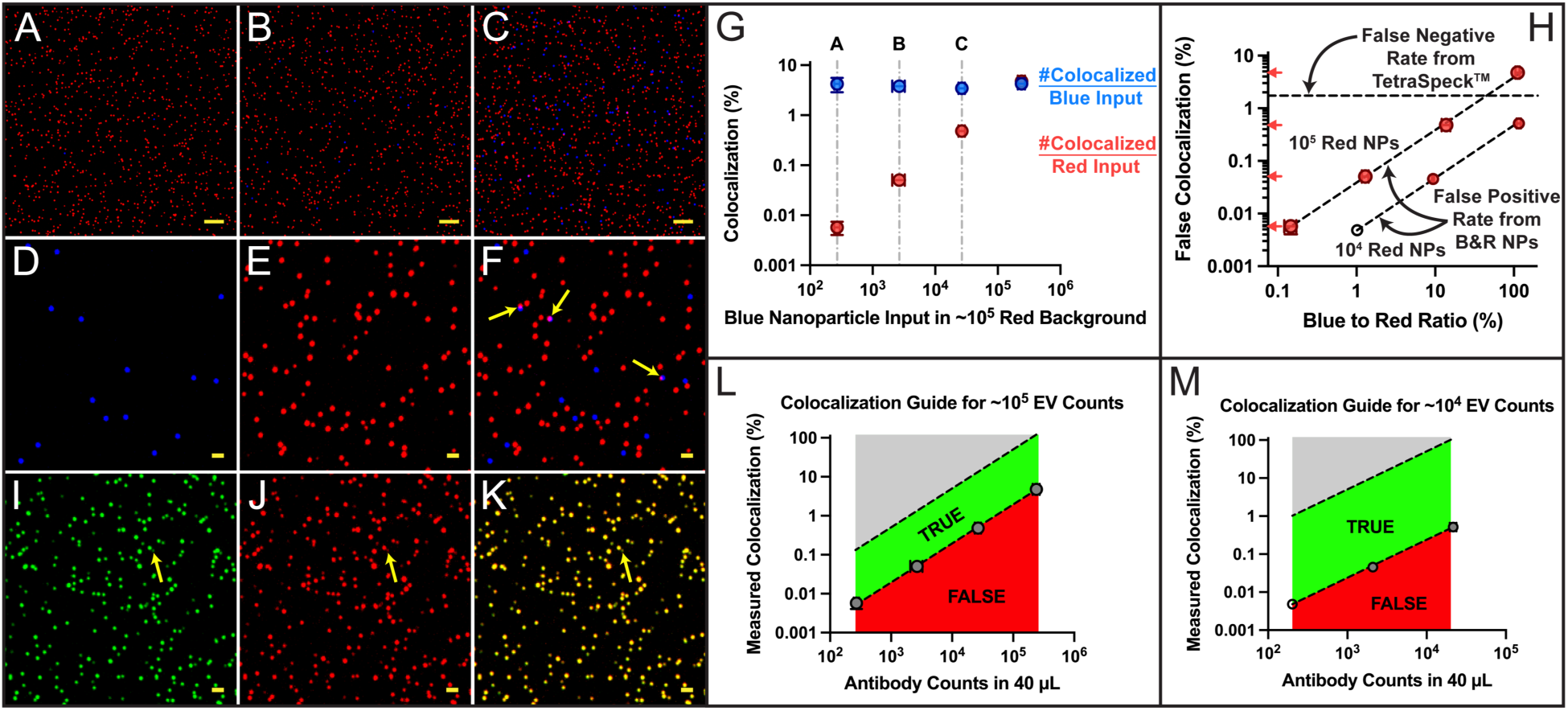
Nanoparticle Colocalization Control Experiments. **(A-F)** False positive colocalization control experiments were completed using two nanoparticle species labeled with distinct fluorescent dyes (red or blue). The red nanoparticle input was held constant (10^5^) and the blue nanoparticle input spanned four orders of magnitude (10^2^ - 10^5^). (A-C) Representative regions of interest (ROI = 119 pm x 119 pm) in confocal microscope images taken following nanoparticle capture on ultrathin nanoporous silicon nitride (NPN) membranes. **(A), (B),** and **(C)** correspond with nanoparticle inputs of 10^5^ red and 10^2^ blue, 10^5^ red and 10^3^ blue, and 10^5^ red and 10^4^ blue, respectively. Scale bars = 10 pm. (D-F) Representative region of interest (ROI = 39 pm x 39 pm) in confocal microscope images taken following capture of 10^5^ red and 10^4^ blue nanoparticles. **(D), (E),** and **(F)** correspond with blue channel signal, red channel signal, and merged signals (blue & red), respectively. Arrows in (F) indicate instances of false positive colocalization detection. Scale bars = 2 pm. (G) Measured colocalization rates are presented relative to both blue and red inputs (indicated by color). The labels A-C denote nanoparticle combinations associated with images shown in panels **(A-C). (H)** False positive colocalization rates relative to red nanoparticle input presented in terms of blue:red nanoparticle ratio (%). Larger data points and tick marks shown on y-axis correspond with false positive colocalization curve completed with 10^5^ red nanoparticles; smaller data points correspond with false positive colocalization curve completed with 10^4^ red nanoparticles. Empty data point on 10^4^ red nanoparticle curve represents a predicted value using data associated with 10% and 100% blue:red nanoparticle input ratios. A single input (10^5^) of TetraSpeck™ fluorescent microspheres labeled with multiple fluorescent dyes was used to determine the false negative colocalization detection rate (presented with dashed horizontal line). (I-K) Representative region of interest (ROI = 39 pm x 39 pm) in confocal microscope images taken following capture of TetraSpeck™ microspheres. **(I), (J),** and **(K)** correspond with green channel signal, red channel signal, and merged signals (green & red), respectively. The arrow in (l-K) indicates an instance of false negative colocalization. Scale bars = 2 pm. (L) Guide for colocalization detection using blue (10^2^ - 10^5^) and red (10^5^) nanoparticles to simulate independent antibody and pan EV signals, respectively. Green and red regions depict putative regions of true and false positive colocalization detection, respectively. (M) Guide for colocalization detection using blue (10^2^ - 10^4^) and red (10^4^) nanoparticles to simulate independent antibody and pan EV signals, respectively. Green and red regions depict putative regions of true and false positive colocalization detection, respectively. Empty data point on curve represents a predicted value calculated as mentioned previously. All captured nanoparticles were imaged with a fluorescence microscope and quantification and colocalization analyses were completed using an open-source FIJI plugin (ComDet v.0.5.5). All data points represent mean ± SEM (n = 3)

We reasoned that when faced with an unacceptably high false positive rate, an investigator could lower the number of pores occupied by EVs to reduce the probability of false colocalization with disassociated antibody labels. To illustrate this benefit we reduced the red nanoparticle background by 10X. We again saw a linear increase in false positives with the blue nanoparticle input but at a 10X lower rate (**Figure 4H**). Thus, if the pan EV signal density is high (>10^5^) and rare colocalization events are of interest, reducing the EV input to 10^4^ would decrease the rate of false positives resulting from the nearby capture of independent signals. Of course, diluting EV samples further increases the scarcity of rare events and may require surveying more FOVs, the entire membrane (**Figure 2C**), or possibly multiple devices. We did not observe any false positive colocalization events with inputs of 10^4^ red and 10^2^ blue nanoparticles, but this is likely due to the limited number of FOVs inspected. Using false positive rates associated with blue:red input ratios of 10% and 100% to predict the behavior of 1%, we anticipate a false positive rate of ∼0.005% which corresponds with one event occurring across 54 FOVs. In our experiments, 15 FOVs (5 FOVs in 3 devices) were analyzed, and thus, our finding of 0% is not surprising.

Based on the above results, we sought to develop a guide to predicting the false positive rate caused by random colocalization as a function of the measured antibody:pan EV signal ratio. At an input of 10^5^ EVs, we predict rates of 0.050%, 0.482%, and 4.759% at antibody:pan EV ratios of ∼1%, ∼10%, and ∼100%, respectively (**Figure 4H**). At an input of 10^4^ EVs, these errors are reduced to 0%, 0.046%, and 0.514%, respectively (**Figure 4H**). We summarize these results in ‘user-friendly’ colocalization guides for EV inputs of 10^5^ (**Figure 4L**) and 10^4^ (**Figure 4M**). Putative regions of true positive (green) and false positive (red) colocalization have been denoted to explicitly define CAD-LB’s capacity for biomarker detection.

We emphasize that the above experiments and analysis only guard against misclassifying the random colocalization of independent signals as specific labeling. EV experiments will require isotype controls to determine non-specific labeling levels which will ultimately be used as correction factors. However, antibody solutions typically contain aggregates, large enough to be captured by NPN pores, and the degree of aggregation may depend on their unique production and purification. Furthermore, antibodies may label soluble forms of proteins targeted on EVs to again create complexes large enough to be captured. The guides presented here provide a specificity metric that incorporates total antibody signal density to avoid misidentifying colocalization that is not EV-associated. Another scenario where metrics like these are useful is biomarker multiplexing; it is possible that two EVs that are positively labeled for unique biomarkers are perceived as a single “double-positive” EV. Depending on the relative abundance of both antibody signals, empirical false positive rates can be used to determine if double labeling is specific or artifactual.

### CAD-LB Quantification of Purified EVs

Having assessed CAD-LB’s performance with well-defined nanoparticles, we next examined performance with EVs using different pan EV labels. First, EVs were collected from the conditioned cell culture media of primary bladder epithelial (PBE) cells *via* conventional ultracentrifugation. PBE EVs were characterized by NTA to establish an approximate stock concentration and a dilution series was used to prepare input samples spanning five orders of magnitude. CAD-LB was performed in the same manner as with nanoparticles (**Figure 5A-C**). Prior to capture, EVs were labeled with one of two pan EV markers: CFSE or MemGlow^TM^ 488 which should label EV luminal proteins and lipid membrane components, respectively. We found that CAD-LB generated a linear response (R^2^ = 0.997) for both CFSE- and MemGlow^TM^-labeled EV samples (**Figure 5D**). As with nanoparticles, however, the highest input concentration tested resulted in images that appear saturated and unreliable for quantification. We find it meaningful that both pan EV labels resulted in highly similar counting curves despite labeling unique EV components (lumen vs. membrane). This finding builds further confidence in CAD-LB, both labels, and the fact that signals are EV-derived. If any one of these factors was compromised, we would not expect to see a strong accordance between CFSE and MemGlow^TM^. Thus for the purposes of counting, a conventionally purified EV sample exhibits a dynamic range comparable to that of nanoparticles (∼10^2^ - 10^6^). Similar to nanoparticle findings, we note that CAD-LB can extend EV dynamic range by at least an order of magnitude compared to alternative single EV technologies employing microscopy^43^ and nFC^21^ techniques.

**Figure 5.**
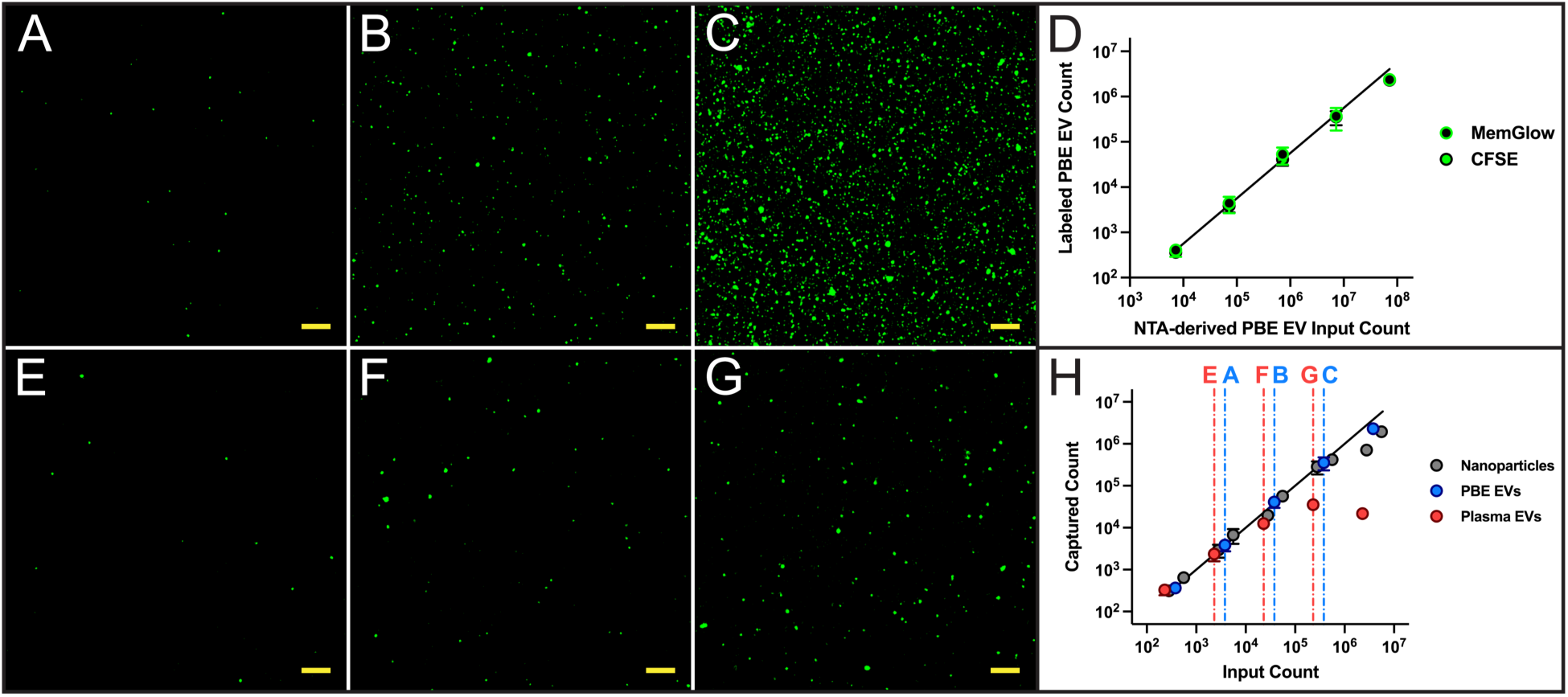
CAD-LB EV Quantification. **(A-C)** Representative regions of interest (ROI = 119 pm x 119 pm) in confocal microscope images taken following purified EV capture. EVs were collected from the conditioned cell culture media of primary bladder epithelial (PBE) cells *via* ultracentrifugation. EVs were characterized with nanoparticle tracking analysis (NTA) for reference, labeled with carboxyfluorescein succinimidyl ester (CFSE), and diluted to prepare inputs spanning five orders of magnitude. **(A), (B),** and **(C)** correspond with EV capture counts of 4-10^3^, 4-10^4^, and 410^5^ EVs, respectively. Scale bars = 10 pm. (D) PBE EVs were labeled with CFSE or MemGlow 488™ and processed with CAD-LB to compare quantification behavior when labeling EV lumen and lipid membrane components, respectively. **(E-G)** Representative regions of interest (ROI = 119 pm x 119 pm) in confocal microscope images taken following plasma EV capture. Plasma samples were labeled with CFSE, prefiltered with SepCon™ spin cups (0.5 pm pores) and diluted to prepare inputs spanning five orders of magnitude. **(E), (F),** and **(G)** correspond with EV capture counts of 2-10^3^, 1-10^4^, and 410^4^ EVs, respectively. Scale bars = 10 pm. **(H)** CAD-LB counting curves for nanoparticles, PBE EVs, and plasma EVs. All counting curves were aligned based on capture counts to compare quantification behavior among different particle sources. The labels **A-C** and **E-G** denote EV capture counts associated with images shown in panels **(A-C)** and **(E-G).** All captured species were imaged with a fluorescence microscope and quantified using an open-source FIJI plugin (ComDet v.0.5.5). All data points represent mean ± SEM (n = 3).

We note that counts of EVs by CAD-LB are roughly an order of magnitude lower than expected based on NTA measurements (**Figure 5D**). However, NTA has been shown to overestimate sample concentration^16–20^ and nFC methods have reported nanoparticle and EV concentrations that are 10-fold and 20-fold less, respectively, than those determined by NTA.^21^ Similarly, both nFC and tRPS have reported EV concentrations roughly 4-fold lower than NTA for identical samples.^18^ Importantly, experiments using liposomes with calculated concentration values (based on size and composition) indicated that nFC and tRPS methods were more accurate compared to NTA.^18^ NTA detects particles based on scattered light which could result in contaminating species, such as protein aggregates, contributing to concentration estimations. The fluorescent dyes used for CAD-LB add specificity to EV quantification since distinct EV components (lumen and membrane) are labeled. While recognized EV quantification methods (NTA, nFC, and tRPS) are used somewhat interchangeably, there is considerable discrepancy among these techniques.^18, 21^ A standardized EV quantification method that offers high degrees of accuracy and throughput would be valuable for the field, but would particularly benefit therapeutic applications.^18, 49^ The therapeutic potential of an EV preparation may be directly related to total EV quantity and a simple tool that provides accurate concentration measurements would prove useful when conducting potency assays.^12^ An EV tool that can provide both accurate concentration and biomarker measurements would further enable therapeutic testing. For example, EV-based tumor vaccination clinical trials have previously used concentrations of major histocompatibility complex (MHC) class II molecules in EV preparations for dosing purposes.^50, 51^

### CAD-LB Quantification of Plasma EVs

Following purified EV studies we assessed the ability of CAD-LB to directly quantify the EVs in plasma. Plasma samples were labeled with CFSE and prefiltered using SepCon^TM^ spin cups containing microporous membranes (0.5 µm pores) (**Figure 2C: Bottom Left Inset**) to remove large aggregates and contaminating debris. Following prefiltration, samples were diluted and input samples spanning five orders of magnitude were processed by CAD-LB (**Figure 5E-G**). For the purpose of comparing quantification behavior among different particle sources, counting curves were aligned based on capture counts (**Figure 5H**). Despite observing comparable linear dynamic ranges for nanoparticle and purified EV samples that extend to inputs of 10^6^, plasma EV samples result in non-linearity at inputs above 10^4^. While the quantification of nanoparticles and purified EVs were constrained by the limits of optical imaging, the 10^4^ value is well below these limits. We believe this early plateau in dynamic range is caused by the presence of contaminating species in plasma such as lipoproteins and protein complexes including aggregates.^52–55^ These EV-sized contaminants could occupy NPN pores and compete with EVs for capture. Despite this decrease in dynamic range for accurate counting, the analytical sensitivity and accuracy of CAD-LB still enable rare biomarker detection with tens of thousands of EVs captured directly from plasma. Preprocessing steps that reduce contaminant levels could increase EV capture density and extend the dynamic range of plasma samples. In summary of the quantification findings, CAD-LB exhibits a wide linear dynamic range: ∼10^2^ - 10^6^ for nanoparticles and purified EVs, which is limited to ∼10^2^ - 10^4^ for plasma-derived EVs.

### CAD-LB Biomarker Validation 1: Detection of CD9 on EVs

We next performed two independent studies to validate the ability of CAD-LB to detect biomarkers on EVs. In the first, we compared CAD-LB and Western blot measurements of CD9 on EVs collected from the conditioned media of mesenchymal stem cells (MSCs) and fibroblasts *via* ultracentrifugation. Transmission electron microscopy (TEM) confirmed vesicle morphology (**Figure S3A**) and NTA estimated particle size and concentration (**Figure S3B**). The tetraspanin proteins (CD9/CD63/CD81) are commonly used in EV analyses since they are expected to be present in EV isolates^56^ and are enriched in EVs relative to cell lysates.^57^ We first confirmed CD9 expression in both MSC and fibroblast EV samples using Western blot. Next, we evaluated CD9 expression in both EV samples using CAD-LB and confirmed that a measurable subset of EVs was CD9+ (**Figure 6A-F**). CFSE was used as the pan EV label (**Figure 6A & 6D**) and a fluorescently-conjugated antibody was used to target CD9 (**Figure 6B & 6E**). CAD-LB successfully detected colocalization of CD9 with EV signals for both MSC and fibroblast samples (arrows in **Figure 6C & 6F**). CD9 labeling was specific since the observed colocalization levels were significantly higher than corresponding isotype controls (**Figure 6G**) and surpassed our false positive rate thresholds (**Figure 4L**) by 15.6-fold and 7.1-fold for MSC and fibroblast samples, respectively. After correcting for isotype control levels, CAD-LB reported that 6.4% of MSC EVs and 0.9% of fibroblast EVs were CD9+ (**Figure 6H**).

**Figure 6.**
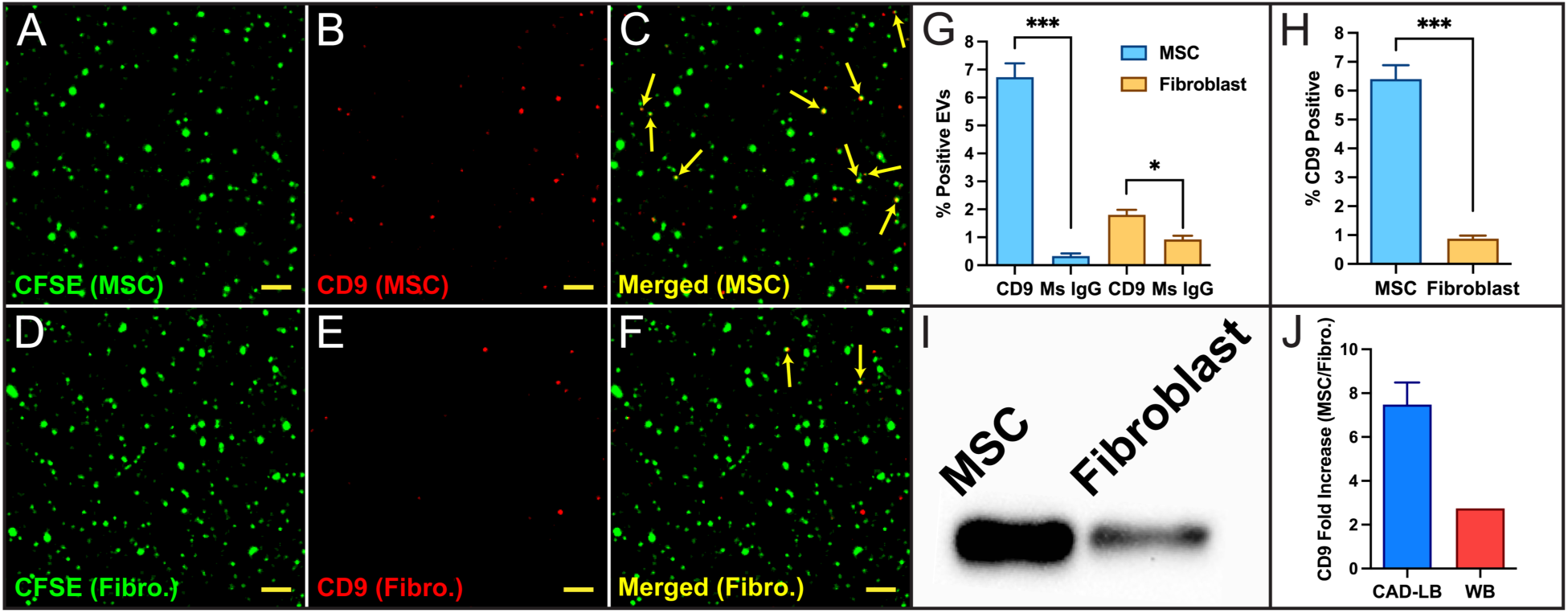
CAD-LB CD9 Detection Validation on EVs. **(A-F)** Representative regions of interest (ROI = 51 pm x 51 pm) in confocal microscope images taken following purified EV capture. EVs were collected from the conditioned cell culture media of mesenchymal stem cells (MSCs) and fibroblasts *via* ultracentrifugation and were evaluated using an antibody targeting CD9 and an appropriate isotype control (mouse [Ms] IgG). EVs were characterized with nanoparticle tracking analysis, labeled with carboxyfluorescein succinimidyl ester (CFSE) then antibodies, prefiltered with SepCon™ spin cups (0.5 pm pores), and diluted to prepare an input of 10^5^ EVs. **(A), (B),** and **(C)** correspond with the CFSE signal, CD9 antibody signal, and merged signals (CFSE & CD9) fora representative MSC EV sample, respectively. **(D), (E),** and (F) correspond with the CFSE signal, CD9 antibody signal, and merged signals (CFSE & CD9) for a representative fibroblast EV sample, respectively. Arrows in **(C)** and **(F)** indicate instances of colocalization and detection of CD9+ EVs. Scale bars = 5 pm. **(G)** Detection summary for CD9 and isotype control in terms of % positive EVs for MSC and fibroblast samples. (H) CD9 detection summary for MSC and fibroblast EVs following isotype control correction. (I) Corresponding Western blot (WB) bulk assay comparison for CD9 expression in MSC and fibroblast EV samples. (J) Comparison of CD9 fold increase (MSC/fibroblast) as reported by CAD-LB and WB assays. All captured EVs were imaged with a fluorescence microscope and quantification and colocalization analyses were completed using an open-source FIJI plugin (ComDet v.0.5.5). All bars represent mean ± SEM (n = 3). * p < 0.05, *** p < 0.001, by Student’s t-test.

The Western blot bulk assay used the same number of EVs from each cell type and detected that MSC EVs were far more enriched in CD9, exhibiting 2.7-fold higher expression, compared to fibroblast EVs (**Figure 6I**). Because CAD-LB returned the same biomarker relationship as Western blot, we consider this test a validation of the ability of CAD-LB to detect relative levels of EV surface proteins among different samples. It is noteworthy that CAD-LB revealed a much higher 7.5-fold difference in CD9 expression between the MSC and fibroblast samples (**Figure 6J**). This disparity is not surprising given the intrinsic differences between a ‘digital’ (CAD-LB) and bulk (Western blot) EV assay. One scenario where this disparity would be observed is if there was differential protein cargo loading between MSC and fibroblast EVs. For instance, if fibroblast EVs are packaged with more CD9 per vesicle, a bulk assay would report a smaller fold enrichment in MSC samples relative to a single EV assay detecting the percentage of CD9+ EVs. Additionally, the discrepancy among assays may in part be attributed to using different detection antibodies designed for unique applications (flow cytometry vs. Western blot).

The different sizes of CD9+ EV subsets identified for MSC and fibroblast samples are consistent with published findings that indicate tetraspanin expression patterns depend on parent cell type.^43, 58–60^ Moreover, there is no ‘gold standard’ technique to compare CAD-LB findings with and current single EV technologies report highly variable CD9+ percentages (∼5 - 89%)^22, 29, 44, 47, 61, 62^ highlighting the impact of both the parent cell type and analysis technique. Other methods report CD9+ percentages but are biased as a result of neglecting tetraspanin-negative EVs^43, 59^ or using capture antibodies that enrich specific subsets.^58, 63^ CAD-LB advantageously captures the entire EV population which should result in an unbiased biomarker assessment.

### CAD-LB Biomarker Validation 2: Detection of ICAM-1 on EVs Secreted by TNF-α Stimulated HUVECs

For our second validation of biomarker detection by CAD-LB, we measured the level of intercellular adhesion molecule 1 (ICAM-1) on EVs secreted by human umbilical vein endothelial cells (HUVECs) stimulated with the inflammatory cytokine tumor necrosis factor alpha (TNF-α). ICAM-1 is known to be significantly upregulated on the surface of vascular endothelial cells under inflammatory conditions^64^ and secreted in soluble form (sICAM-1) upon activation.^65, 66^ Recently a fraction of the sICAM-1 has been shown to be EV-associated,^67^ a result that should be readily confirmed with CAD-LB.

In these experiments, HUVEC cells were either stimulated with TNF-α or left untreated for 24 hours. HUVEC cells were then stained for ICAM-1 expression using immunocytochemistry (ICC) methods (**Figure 7A-C**) and EVs from the conditioned cell culture media were evaluated for ICAM-1 expression *via* CAD-LB (**Figure 7D-F**). As expected, TNF-α stimulated HUVECs exhibited a substantial upregulation of surface ICAM-1; ICC revealed an approximate 37-fold increase in mean fluorescence intensity (MFI) (**Figure 7C**). Concomitantly, CAD-LB successfully detected ICAM-1+ EV subsets in the conditioned media of both stimulated and unstimulated HUVECs (**Figure 7D-E**). Indeed, CAD-LB returned the same ICAM-1 expression relationship as ICC identifying 28.3% and 4.0% ICAM-1+ EVs in stimulated and unstimulated HUVEC conditioned media samples, respectively (**Figure 7F**). Compared to cellular expression changes, the EV compartment was altered to a lesser degree with stimulated HUVEC-derived EVs presenting 7.5-fold higher ICAM-1 expression. Importantly, isotype controls returned colocalization levels orders of magnitude lower for both stimulated and unstimulated samples (**Figure 7G**: representative stimulated sample), and ICAM-1 colocalization rates were at least an order of magnitude above the thresholds for false positives originating from random capture (**Figure 4M**).

**Figure 7.**
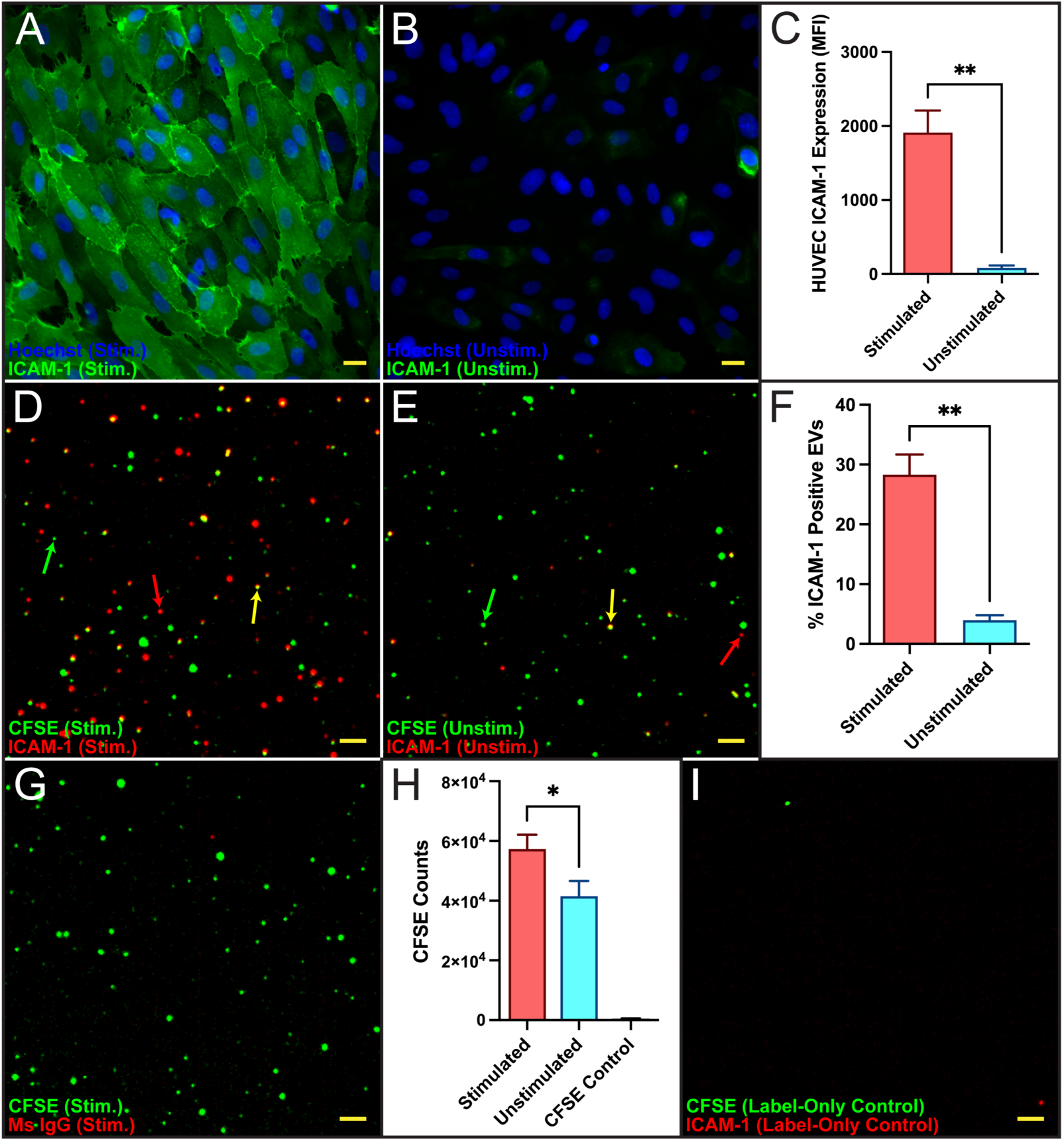
CAD-LB ICAM-1 Detection on EVs Secreted by TNF-a Stimulated HUVECs. **(A-B)** Representative epifluorescence microscope images (333 µm x 333 µm) showing merged signals (Hoechst & ICAM-1) taken following HUVEC treatment. Stimulated (TNF-a) **(A)** and unstimulated **(B)** HUVECs were stained for ICAM-1 expression using ICC methods. Scale bars = 20 µm. **(C)** ICAM-1 mean fluorescence intensity (MFI) summary for HUVEC samples following background correction (secondary antibody-only). **(D-E)** Representative regions of interest (ROI = 71 µm x 71 µm) in confocal microscope images taken following EV capture. The conditioned media of HUVECs was directly evaluated for the presence of ICAM-1 + EVs *via* CAD-LB. Conditioned media was labeled with carboxyfluorescein succinimidyl ester (CFSE) then antibodies, prefiltered with SepCon™ spin cups (0.5 µm pores) and diluted to prepare an input of 104 EVs. **(D)** and **(E)** show the merged signals (CFSE & ICAM-1) for a representative stimulated and unstimulated HUVEC conditioned media sample, respectively. The green, red, and yellow arrows in **(D)** and **(E)** indicate a lone CFSE signal, a lone ICAM-1 signal, and an instance of colocalization, respectively. Scale bars= 5 µm. **(F)** ICAM-1 detection summary for HUVEC-derived EVs following isotype control correction. **(G)** Region of interest (ROI= 71 µm x 71 µm) showing merged signals (CFSE & mouse [Ms] lgG isotype control) in representative stimulated HUVEC conditioned media sample. Scale bar = 5 µm. **(H)** CFSE count summary for stimulated and unstimulated HUVEC conditioned media samples (n = 6) and label-only (no EV) controls. **(I)** Region of interest (ROI= 71 µm x 71 µm) showing merged signals (CFSE & ICAM-1) in representative label-only (no EV) control sample. Scale bar = 5 µm. EV quantification and colocalization analyses were completed using an open-source FIJI plugin (ComDet v.0.5.5). Unless otherwise indicated, all bars represent mean± SEM (n = 3). * p < 0.05, ** p < 0.01, by Student’s t-test.

The finding that ICAM-1 expression is upregulated on HUVEC EVs following TNF-α stimulation is a phenomenon previously reported and carefully verified by Hosseinkhani *et al.*^67–69^ Interestingly, EVs derived from stimulated HUVECs are known to act as a functional mediator between endothelial cells and monocytes.^68^ Hosseinkhani *et al.* uncovered that TNF-α stimulation triggers the release of EVs capable of upregulating recipient endothelial cell ICAM-1 expression and inducing monocyte migration. In further accordance with published results,^67^ CAD-LB detected a higher concentration of EVs when HUVECs were stimulated with TNF-α (5.7⋅10^4^ vs. 4.1⋅10^4^ captured EVs, *p < 0.05*) (**Figure 7H**). Both CAD-LB and NTA^67^ report a nearly identical fold increase in EV concentration following TNF-α stimulation (1.4-fold and 1.3-fold, respectively). Crucially, both CAD-LB experimental groups yielded CFSE counts orders of magnitude higher than CFSE-only controls (**Figure 7H-I**). Of note, proteolytic cleavage is known to generate sICAM- 1 following endothelial cell activation.^65^ A fraction of the total ICAM-1 CAD-LB signal may be derived from free sICAM-1 protein, however, EV biomarker detection is contingent upon colocalization with CFSE. We did observe ICAM-1 signal unassociated with CFSE, and the unassociated fraction was larger for each stimulated replicate compared to their unstimulated counterparts. Thus, lone ICAM-1 signal may in part represent the specific labeling of free sICAM- 1.

The HUVEC TNF-α stimulation experimental model revealed that CAD-LB successfully detected EV biomarkers that capture protein expression changes in parent cells. Additionally, the ability of CAD-LB to discriminate EV protein profiles that reflect cellular functional states strengthens its position as a liquid biopsy assay. Not only do these findings further validate CAD-LB biomarker detection, but they also illustrate the potential for diagnostic utility. Namely, EV-associated ICAM-1 may have value in identifying individuals with coronary heart disease.^67^ One important operational detail in these experiments is that CAD-LB ICAM-1 detection was completed with conditioned cell culture media cleared only of cellular debris. This highlights CAD-LB’s capacity to process samples without EV enrichment owed to low EV input requirements and the ability to pass high concentrations of contaminating protein species.^45^ This direct labeling model obviates complex EV isolation protocols, which can each uniquely bias downstream analyses^70, 71^ and yield low-purity EV samples,^55, 72^ and ultimately demonstrates the rapid and simple nature of CAD-LB.

### EV Biomarker Interrogation in Patient Samples

To demonstrate the clinical potential of CAD- LB biomarker detection, plasma EVs were evaluated for the presence of diagnostically valuable biomarkers. Immune checkpoint blockade (ICB) is a promising cancer immunotherapy that has revolutionized the treatment of various malignancies.^73, 74^ ICB targets cytotoxic T-lymphocyte associated protein 4 (CTLA-4) and programmed cell death protein/ligand 1 (PD-1/PD-L1) immunosuppressive signaling pathways to reinvigorate and sustain T-cell-mediated anti-tumor responses. PD-L1 is expressed by various cells in the tumor microenvironment (TME) as well as tumor-derived EVs that contribute to immune evasion.^75–77^ Chen *et al.* reported that EV-associated PD-L1 could be used to identify treatment responders among melanoma patients receiving α-PD- 1 ICB.^75^ For this reason, we tested the feasibility of detecting PD-1 and PD-L1 biomarkers in patient samples using CAD-LB.

Levels of EV-associated PD-1 and PD-L1 were assessed in plasma collected from a healthy donor, a bladder cancer patient who had undergone α-PD-L1 ICB (atezolizumab), and a bladder cancer patient who had undergone α-PD-1 ICB (pembrolizumab). CAD-LB was used to process plasma samples and determine the percentages of biomarker-positive EVs (**Figure 8**). The findings presented here are promising and suggest that even rare biomarkers (<1%) with diagnostic value can be detected above isotype control levels. Additionally, it is worth noting that EV PD-L1 levels are higher than EV PD-1 levels for all individuals observed which is encouraging since more cell types are known to express the former.^78^ The predictive power of biomarkers detected by CAD-LB cannot be assessed at this stage since sampling timepoints and clinical outcomes are unknown. However, since the detection of clinically useful biomarkers appears feasible, replicating this study with a larger donor cohort, increased sampling timepoints and known treatment responses is our next goal. We also note that while a healthy donor control was included in these experiments, it is not meaningful to make any comparisons to ICB patients since each experimental group only includes technical replicates from single donors and is not representative of matched control and treatment groups.

**Figure 8.**
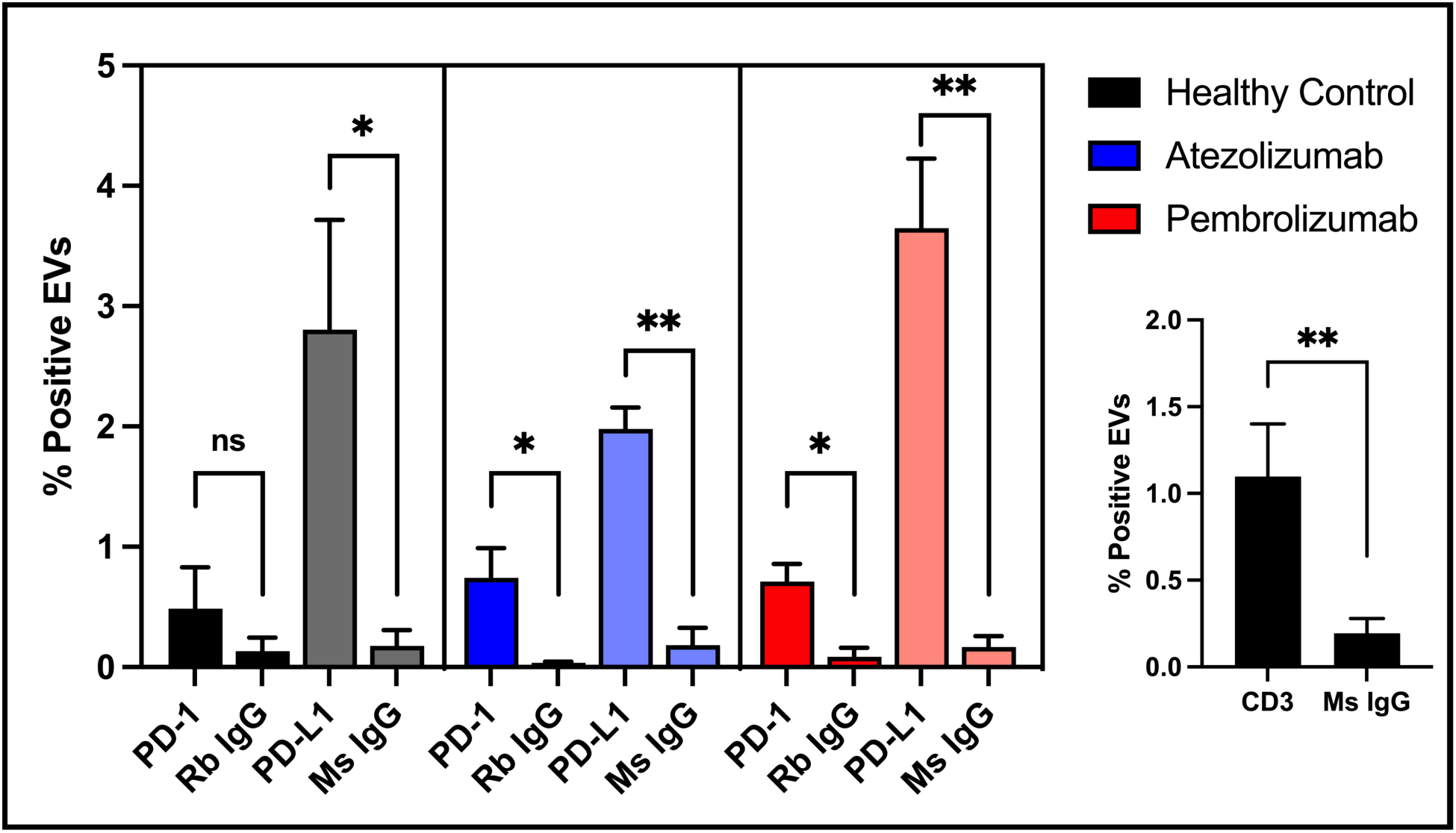
CAD-LB EV Biomarker Detection in Healthy Control and Bladder Cancer Patient Plasma Samples. Healthy donor control and bladder cancer patient plasma samples were evaluated using antibodies targeting PD-1 and PD-L1 (or CD3) and their respective isotype controls, rabbit (Rb) and mouse (Ms) IgG antibodies. Samples were labeled with carboxyfluorescein succinimidyl ester (CFSE) then antibodies, prefiltered with SepCon™ spin cups (0.5 pm pores) and diluted to prepare an input of 10^5^ EVs. Patient samples are categorized by treatments received: **atezolizumab** (a-PD-LI ICB) and **pembrolizumab** (o-PD-1 ICB). All captured EVs were imaged with a fluorescence microscope and quantification and colocalization analyses were completed using an open-source FIJI plugin (ComDet v.0.5.5). All groups represent technical replicates from single donors. All bars represent mean ± SEM (n = 3). * p < 0.05, ** p < 0.01, ns : not significant, by Student’s t-test.

An important extension of CAD-LB, beyond the foundational work presented here, is its potential to interrogate multiple biomarkers simultaneously. Biomarker multiplexing would enable the identification of EV parent cells as well as biomarkers of disease or therapeutic response. The potential for immune cell assessment was illustrated through the successful detection of the T cell marker CD3 by CAD-LB (**Figure 8: Inset**). Multiplexing PD-1 (and complementary functional biomarkers) with CD3 could elucidate the relative proportions of various T cell functional states, including those that are activated and exhausted. These metrics may help to reveal the composition of TMEs and lymphoid tissues and provide a means for monitoring the systemic immune response. In doing so, we would establish a noninvasive method for monitoring the efficacy of immunotherapies, like ICB, in real-time. These ambitions appear to be running into the current limits of CAD-LB for single EV detection above isotype control levels. Specifically, <1% of plasma EVs were CD3+, and similarly, <1% were PD-1+. Thus, the fraction of CD3+ EVs that are also PD-1+ can be expected to be <0.01%. This means that detailed analyses of CD3+ EV subsets are likely unfeasible and would require enrichment of target cell EVs (anti-CD3 affinity purification) upstream of CAD-LB. Additional technical advances in detection and isotype control antibodies, possibly combined with imaging system improvements, may help minimize sample preprocessing to retain the simplicity of CAD-LB for biomarker detection.

## CONCLUSION

The interest in EVs has grown at a rapid pace due to their potential in diagnostic and therapeutic applications. In parallel, the development of EV-based technology has also accelerated. Bulk EV analyses cannot capture EV heterogeneity and are inadequate for studying rare subsets relevant to disease. While sensitive techniques that can analyze single EVs have been developed, most suffer from complicated protocols, prohibitive instrumentation, limited throughput, and the inability to multiplex biomarkers. In this work, we rigorously characterized and validated CAD-LB as a microfluidic strategy that enables the rapid isolation of EVs and provides a tool for EV quantification and surface protein biomarker assessment. CAD-LB enables wide adoption due to its simplistic operation and minimal instrumentation. Our findings provide evidence that CAD-LB accurately measures EV biomarkers which can reflect parent cell status. We believe CAD-LB is a useful platform well suited for monitoring valuable EV biomarkers in various settings.

## EXPERIMENTAL SECTION

### µSiM Device Assembly

µSiM components and assembly jigs were purchased from ALine Inc. (Signal Hill, CA.). Ultrathin nanoporous membranes were purchased from SiMPore Inc. (West Henrietta, NY; Nanoporous Membrane, **Cat. # NPSN100-1L, NPSN100-2L, NPSN100-2LWM**). NPN membrane fabrication^32, 79^ yields a randomly distributed pore array with an average pore size of ∼60 nm. µSiM device assembly has been described in detail previously.^38^ Briefly, µSiM devices were assembled by applying pressure to membranes and acrylic components lined with pressure-sensitive adhesive; components were aligned using assembly jigs.

### Tangential Flow for Analyte Capture (TFAC) Device Fabrication

Custom polydimethylsiloxane (PDMS) sheets (100 & 300 μm) (Trelleborg Sealing Solutions Americas, Fort Wayne, IN) were patterned using a Silhouette Cameo digital craft cutter (Silhouette America, Oren, UT) as described previously.^35^ Patterned silicone components were stacked layer-by-layer to assemble the TFAC microfluidic device. NPN membranes were situated within silicone layers such that microfluidic channels were established above and beneath the membrane. The 100 μm top channel was UV/ozone treated and boned to a PDMS support component in a 70°C oven. The PDMS/top channel component, remaining silicone layers (300 μm), NPN membrane, and a glass slide base were sealed for flow using a custom-machined aluminum clamp.

### Nanoparticle Sample Preparation

Fluorescent polystyrene nanoparticles were used to assess CAD-LB’s ability to quantify nanoparticles and detect colocalization events. Nanoparticles labeled with a single fluorescent dye (Thermo Fisher Scientific, **Cat. # B150**, **G100**, **R100**) were used for nanoparticle quantification (G100) and false positive colocalization detection (B150 & R100). Nanoparticles were provided at a known concentration (1.8⋅10^13^ nanoparticles/mL [G100 & R100] or 5.3⋅10^12^ nanoparticles/mL [B150]), calculated using the formula: (6⋅10^10^⋅S)/(π⋅D^3^⋅ρ), where S is weight % solids (1%), D is mean nanoparticle diameter, and ρ is polymer density (polystyrene ∼1.06 g/cm^3^). TetraSpeck^TM^ microspheres (Thermo Fisher Scientific, **Cat. # T7279**) labeled with four fluorescent dyes were used for false negative colocalization detection. TetraSpeck^TM^ microspheres were provided at a known concentration (1.8⋅10^11^ nanoparticles/mL) reported by the manufacturer. Stock nanoparticle solutions were diluted with 1X phosphate buffered saline (PBS) (Thermo Fisher Scientific, **Cat. # 10010023**) to generate desired nanoparticle inputs.

### Cell Culture & EV Sample Preparation

The EVs analyzed in this study were collected from several sources. First, EVs were collected from the conditioned cell culture media of primary bladder epithelial (PBE) cells (ATCC, **Cat. # PCS-420-010**) and human urinary bladder carcinoma grade II (5637) cells (ATCC, **Cat. # HTB-9**). PBE and 5637 cells were cultured using media prepared according to ATCC guidelines and the conditioned cell culture media was collected thrice weekly (during media exchanges) over a 3-month period. Five T75 flasks and two 1000 mL CELLine^TM^ Bioreactor Flasks (Thermo Fisher Scientific, **Cat. # 02-912-742**) were used to maintain PBE and 5637 cells, respectively. EVs were isolated using a Beckman Coulter Optima^TM^ MAX ultracentrifuge (100,000⋅g, 90 minutes) and resuspended in PBS. The PBE sample was characterized with a Malvern NanoSight NS300 NTA instrument to determine an approximate stock solution concentration (∼1.8⋅10^10^ EVs/mL). This concentration was used as reference and PBE EVs were diluted in PBS to obtain desired inputs.

Human mesenchymal stem cells (MSCs; ATCC, **Cat. # PCS-500-012**) were cultured using mesenchymal stem cell basal media with 10% exosome-depleted FBS (Bio-Techne, **Cat. # S11150**; supernatant collected after centrifuging at 100,000⋅g for 18 hours) and 1% Antibiotic-Antimycotic (100X) (Thermo Fisher Scientific, **Cat. # 15-240-062**). Human dermal fibroblasts (ZenBio, **Cat. # DF-F**) were cultured in low glucose Dulbecco’s Modified Eagle’s Medium (Thermo Fisher Scientific, **Cat. # SH30021FS**) with 10% exosome-depleted FBS (Bio-Techne, **Cat. # S11150**; supernatant collected after centrifuging at 100,000⋅g for 18 hours) and 1% Antibiotic-Antimycotic (100X) (Thermo Fisher Scientific, **Cat. # 15-240-062**). Cells were added at 5,000 cells/cm^2^ to 150 mm Petri dishes and media supplemented with a final concentration of 50 µM ascorbic acid 2-phosphate (Sigma Aldrich, **Cat. # A8960**). Media was exchanged every 3 days and cells were cultured for 2.5 weeks; the last 3 days of culture, media was saved for EV isolation. EVs were collected from conditioned cell culture media *via* differential centrifugation (500⋅g, 10 minutes; 2,500⋅g, 20 minutes; and 10,000⋅g, 30 minutes). The resulting supernatant was filtered (0.22 µm) and EVs were isolated using a Beckman Coulter Optima^TM^ ultracentrifuge (100,000⋅g, 70 minutes). EVs were resuspended in 200 µL of particle-free PBS. Isolated EVs were then characterized by transmission electron microscopy (TEM) and NTA. EVs were pipetted onto coated copper grids with a 1% uranyl acetate stain and imaged using a JOEL 1400 TEM with a side mount AMT 2k digital camera to assess morphology. A Malvern NanoSight NS300 NTA instrument was used to determine the average size and concentration of EVs diluted in particle-free water. NTA determined that the stock MSC and fibroblast EV solution concentration was ∼1.8⋅10^10^ EVs/mL and ∼1.2⋅10^10^ EVs/mL, respectively. These concentrations were used as reference and EVs were diluted in PBS to obtain desired inputs.

Human umbilical vein endothelial cells (HUVECs; VEC Technologies, Rensselaer, NY) were cultured using EBM^TM^-2 Endothelial Cell Growth Basal Medium (Lonza Bioscience, **Cat. # CC- 3156**) supplemented with the EGM^TM^-2 Endothelial SingleQuots^TM^ Kit (Lonza Bioscience, **Cat. # CC-4176**) except for the provided FBS supplement. Instead, EBM^TM^-2 basal media was supplemented with 2% exosome-depleted FBS (Thermo Fisher Scientific, **Cat. # A2720803**). Media was exchanged every 24-48 hours and cells were used between passages 3-6. After HUVECs reached confluency, one T25 tissue culture flask was given media spiked with 1 ng/mL recombinant human TNF-α (R&D Systems, **Cat. # 210-TA**) (stimulated group) while another T25 tissue culture flask was only given fresh media (unstimulated group). The conditioned cell culture media from both groups was collected after 24 hours, cleared of cellular components/debris *via* centrifugation (2,000⋅g, 15 minutes), and used for subsequent EV analyses.

EVs were analyzed from the plasma of a healthy donor as well as bladder cancer patient samples purchased from Discovery Life Sciences (Huntsville, AL). Whole blood was collected from a healthy donor following a protocol approved by the University of Rochester Institutional Review Board (IRB). Informed consent was obtained from the healthy donor before blood collection. Whole blood was collected in 10 mL sodium heparin-coated tubes (BD, **Cat. # 366480**) and centrifuged (2,000⋅g, 15 minutes) to generate plasma. Plasma samples were diluted using PBS under the assumption that the initial EV concentration was ∼10^9^ EVs/mL.

### EV Sample Labeling

Collected EV samples were diluted in PBS (except for HUVEC conditioned media) to obtain a 100 µL solution. EVs were labeled with the pan EV label carboxyfluorescein succinimidyl ester (CFSE) (BD Biosciences, **Cat. # 565082**) at a concentration of 5 µM and incubated for 15 minutes in a 37°C water bath. MemGlow^TM^ 488 (Cytoskeleton, **Cat. # MG01**) served as an additional pan EV label in PBE EV quantification experiments; EVs were labeled at a concentration of 200 nM and samples were incubated for 30 minutes at room temperature following the published protocol by Hyenne *et al.*^80^ For biomarker experiments, EVs were labeled with conjugated antibodies: anti-CD9 (Thermo Fisher Scientific, **Cat. # 17-0098-42**), anti-ICAM-1 (Thermo Fisher Scientific, **Cat. # 17-0549-42**), anti-PD-1 (Abcam, **Cat. # ab201825**), anit-PD-L1 (R&D Systems, **Cat. # FAB1561P**), or anti-CD3 (Thermo Fisher Scientific, **Cat. # 12-0038-42**) and corresponding isotype control antibodies: mouse IgG (Thermo Fisher Scientific, **Cat. # 17- 4714-42, MA5-18168**) or rabbit IgG (R&D Systems, **Cat. # IC1051R**) at a concentration of 200 ng/mL and incubated with gentle mixing for 45 minutes at room temperature in the dark. Prior to antibody labeling, HUVEC samples were treated with an Fc receptor blocking reagent (1 test) (Thermo Fisher Scientific, **Cat. # 14-9161-73**) and incubated at 4°C for 15 minutes. Labeled 5637, MSC, fibroblast, HUVEC, and plasma EV samples were subject to microporous membrane (0.5 µm pores) (SiMPore Inc., West Henrietta, NY, **Cat. # MPSN400-3L-0.5HP**) prefiltration using SepCon^TM^ spin cups (SiMPore, Inc., West Henrietta, NY, **Cat. # SC400508**) and filtrate solutions were used for downstream processing. Labeled EV solutions were diluted with PBS to obtain the desired input in 40 µL. For TFAC experiments, EVs were labeled in an identical manner in 1 mL sample volumes. Before TFAC capture, EV samples were diluted to obtain the desired input in 500 µL.

### Western Blot Analysis

3⋅10^8^ total particles were loaded into pre-cast gels (BioRad, **Cat. # 4561094**). Samples were blocked in a 5% nonfat milk solution and stained for the common EV marker CD9 using a primary antibody (Cell Signaling Technologies, **Cat. # 13174**) and an anti-rabbit HRP secondary antibody (Cell Signaling Technologies, **Cat. # 7074**). Samples were imaged using the BioRad Chemidoc chemiluminescent system. FIJI was used to quantify the intensity of bands in Western blots to determine CD9 expression fold difference among EV samples.

### EV Capture in the µSiM

The NPN membrane separates a well structure and microfluidic channel within the µSiM device. Before sample injection, the bottom channel was pre-wet with PBS, and the well was filled with 100 µL of PBS. In experiments where EVs were labeled with conjugated antibodies, the bottom channel was pre-wet with a solution of bovine serum albumin (1 mg/mL in PBS) (Sigma Aldrich, **Cat. # A7906**) which was allowed to incubate for 15 minutes to minimize nonspecific labeling. After the device was primed, double-sided tape (3M^TM^, **Cat. # 468MPF**) was placed over one access port of the microfluidic channel to maintain a dead-end-tangential compound filtration flow scheme. 40 µL of sample was injected *via* pipette into the open access port at a slow, controlled rate.

### EV Capture in TFAC Devices

Flow rates were carefully maintained using syringe pumps connected through microfluidic tubing (1/32” ID silicone tubing; ColeParmer, **Cat. # 95802-01**) to the PDMS support component. The entire fluidic circuit was primed with PBS before flow was initiated. As described previously, the flow was split at the membrane such that a tangential component passed over the membrane while the other fraction was filtered through the membrane.^35, 36^ A supply rate (Q_s_) of 25 µL/min and ultrafiltration rate (Q_u_) of 12.5 µL/min was maintained to process 1 mL samples. Following capture, devices were disassembled and NPN membranes were retrieved and placed inverted in Nunc™ glass bottom dishes (Thermo Fisher Scientific, **Cat. # 150680**) filled with PBS for imaging.

### Immunocytochemistry (ICC)

HUVECs were maintained as described above. For ICC experiments, HUVECs were seeded in tissue culture-treated 12-well plates coated with human fibronectin (5 µg/cm^2^) (R&D Systems, **Cat. # 1918-FN**). Stimulated and unstimulated groups were formed as described above, however, after 24 hours the conditioned cell culture media was discarded and cells were live stained for ICAM-1 expression. HUVECs were washed with media twice and labeled using a mouse primary antibody targeting human ICAM-1 (Thermo Fisher Scientific, **Cat. # 14-0549-82**) at an appropriate working concentration (1:100 dilution in media) and incubated for 15 minutes at 37°C. HUVECs were then washed with PBS, fixed with 4% paraformaldehyde for 10 minutes at room temperature, and washed 3 times with PBS. HUVECs were next blocked with 5% goat serum (Thermo Fisher Scientific, **Cat. # 50062Z**) (diluted in PBS) for 30 minutes at room temperature. HUVECs were then labeled using a goat anti-mouse IgG Alexa Fluor^TM^ 488 secondary antibody (Thermo Fisher Scientific, **Cat. # A-11001**) at an appropriate working concentration (1:200 dilution in PBS) and incubated for 1 hour at room temperature in the dark. HUVECs were washed 3 times in PBS and subsequently labeled with Hoechst (Thermo Fisher Scientific, **Cat. # H3570**) at an appropriate working concentration (1:10,000 dilution in PBS) before imaging. A control well for each cell group included only the secondary antibody and was used for background correction when determining ICAM-1 mean fluorescence intensity (MFI).

### Fluorescence Microscopy

Following the capture process, fluorescence imaging was performed within µSiM devices. Imaging was carried out using an Andor Dragonfly spinning disk confocal microscope equipped with a Zyla 4.2 sCMOS/Sona 2.0B-11 sCMOS camera. A 60X/1.2 NA objective was used to capture images. 405, 488, 561, and 637 nm excitation lasers were used in coordination with 450-50, 525-50, 600-50, and 700-75 nm emission filters, respectively, to visualize fluorescent nanoparticles, pan EV labels, and fluorescent antibodies. 3-5 images (206 µm x 206 µm/238 µm x 238 µm) were taken from each device and used for subsequent analyses. To acquire whole membrane images, the montage and FusionSticher features within the Fusion imaging software were used. To provide a realistic estimate of imaging resolution full width at half maximum intensity values of pseudo point emitters (100 nm fluorescent nanoparticles) were measured in confocal images. For HUVEC ICAM-1 ICC, fluorescence imaging was performed within tissue culture-treated 12-well plates. Imaging was carried out using a Nikon epifluorescence microscope equipped with a Zyla 4.2 sCMOS camera. A 40X/0.55 NA objective was used to capture images. 405 and 490 nm excitation LEDs were used in coordination with 434-32 and 519- 26 nm emission filters, respectively, to visualize Hoechst and ICAM-1 fluorescent labels. 3 images (333 µm x 333 µm) were taken from each well and used for subsequent analyses.

### Quantification & Colocalization Analysis

Confocal fluorescence images were imported into FIJI for quantification and colocalization analysis. An open-source FIJI plugin, ComDet v.0.5.5, was used to identify fluorescence signals and detect instances of signal colocalization among different fluorescence channels. For signal identification, plugin settings used were approximate particle size (pixels) and intensity threshold (SNR). For colocalization detection, the additional plugin setting used was maximum distance between colocalized spots (pixels). The default plugin settings used were 3 (particle size), 5 (intensity threshold), and 2.5 (maximum distance between colocalized spots). Optimal plugin settings were identified for experimental triplicates and maintained for experimental controls. Analysis was completed on all images from the same device and counts were averaged and then extrapolated based on total membrane area. Statistical analysis for quantitative findings was performed using Prism (GraphPad Software, San Diego, CA).

## ASSOCIATED CONTENT

Supporting Information: Figure S1. Nanoparticle quantification systematic error and accompanying theoretical input correction; Figure S2. Nanoparticle dynamic range comparison between CAD-LB and NTA techniques; Figure S3. Example characterization of EVs used for biomarker validation experiments (PDF)

## AUTHOR INFORMATION

### Author Contributions

Conceptualization: J.L.M, J.F., S.N.W., K.L.; Data Acquisition: S.N.W., K.L., M.J.D.; Data Analysis: S.N.W.; Funding Acquisition: J.L.M., J.F.; Project Administration: J.L.M., J.F., S.B., G.H.; Manuscript Writing: S.N.W., J.L.M., J.F.

## ACKNOWLEDGEMENTS

This work was funded by the NIH (R01 EB031581, TL1 TR001858) and DOD (CA170373), as well as the University of Rochester (UR) Technology Development Fund (TDF) and UR Wilmot Cancer Institute (WCI) Shark Tank Pilot program. J.L.M. and J.F. were supported by NIH R01 EB031581 and DOD CA170373. S.N.W. was supported by NIH R01 EB031581, DOD CA170373, and both the UR TDF and UR WCI Shark Tank Pilot program. K.L. was supported by DOD CA170373. M.J.D. was supported by the Clinical and Translational Science Fellowship NIH TL1 TR001858. The authors would like to acknowledge the Center for Biologic Imaging (University of Pittsburgh) for funding used to acquire TEM images. The authors would like to acknowledge Julie Kuebel for their effort toward HUVEC culture maintenance and ICC experiments. We would also like to acknowledge Michael Klaczko for helpful experimental insight and Isabelle Linares for providing HUVECs for stimulation experiments.

