## Supplementary material for "Rapid Assessment of Biomarkers on Single Extracellular Vesicles Using ‘Catch and Display’ on Ultrathin Nanoporous Silicon Nitride Membranes": Figures S1-S3

**This file includes:** Figures S1 - S3

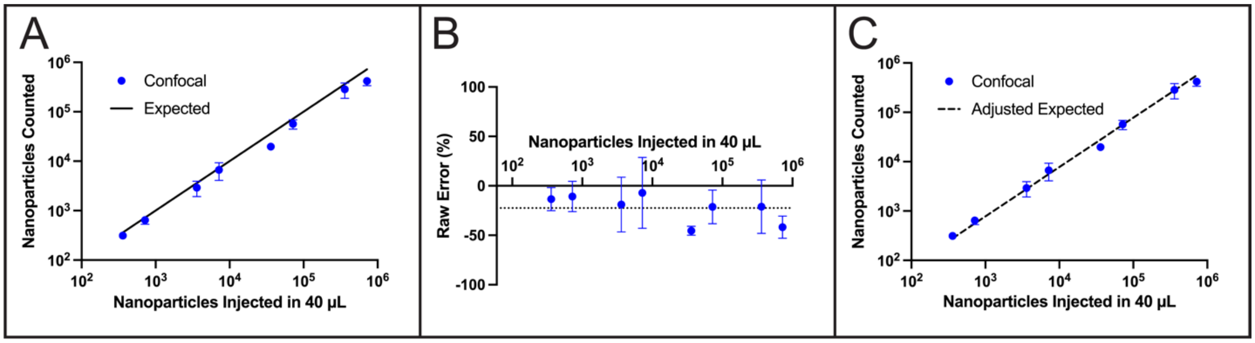


**Figure S1. Nanoparticle Quantification Systematic Error & Correction. (A)** Comparison of nanoparticles counted and theoretical nanoparticle input injected in CAD-LB devices within the linear range of detection. **(B)** Raw percent error within the linear range of detection. Dotted horizontal line indicates average percent error calculated across linear range (-22.5%). **(C)** Comparison of nanoparticles counted and theoretical nanoparticle input using average percent error as correction factor for expected count. All data points represent mean ± SEM (n = 3).

The counting curve presented in **Figure 3** has been adjusted for a systematic error observed in nanoparticle capture experiments (**Figure S1**). The raw error observed across the linear quantification range was consistent and therefore used as a correction factor to adjust all theoretical nanoparticle inputs. CAD-LB undercounts based on the manufacturer's determined concentration which could represent unique biases among quantification methods and/or nanoparticle loss associated with CAD-LB processing. Sample filtration is driven by transmembrane pressure (TMP) provided *via* pipette injection, although when the pipette is removed, TMP is released. To address a concern that nanoparticles would detach following TMP release, live capture videos were taken to assess nanoparticle retention. A region of interest was observed during sample injection, nanoparticle capture, and following TMP release. The density of nanoparticles remained stable following pressure release, and thus, the discrepancy in nanoparticle counts is not due to nanoparticle loss following capture. Additionally, nanoparticle "breakthrough" was assessed by extracting and processing filtrate solutions in secondary devices. Nanoparticle breakthrough counts were <0.1% (data not shown) relative to original capture counts suggesting that breakthrough is not responsible for quantification discrepancy. It is possible that loss occurs upstream of nanoparticle capture during preparation which is likely unavoidable. Furthermore, following the injection of samples, dead volume (~5 µL) exists in the microfluidic channel upstream of the membrane, presenting an opportunity for loss and undercounting.

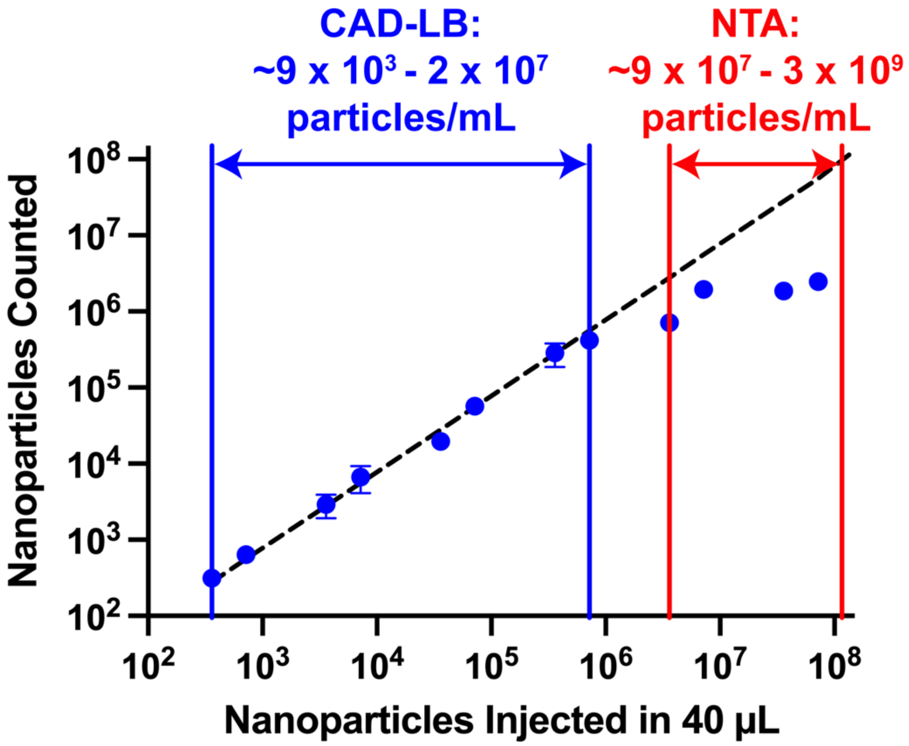


**Figure S2. CAD-LB & NTA Dynamic Range Comparison.** Blue data points: Comparison of nanoparticles counted and theoretical nanoparticle input injected in CAD-LB devices. The approximate dynamic range of CAD-LB and nanoparticle tracking analysis (NTA)[1] assays are shown with blue and red ranges, respectively, in terms of nanoparticle input concentration. All data points represent mean ± SEM (n = 3).

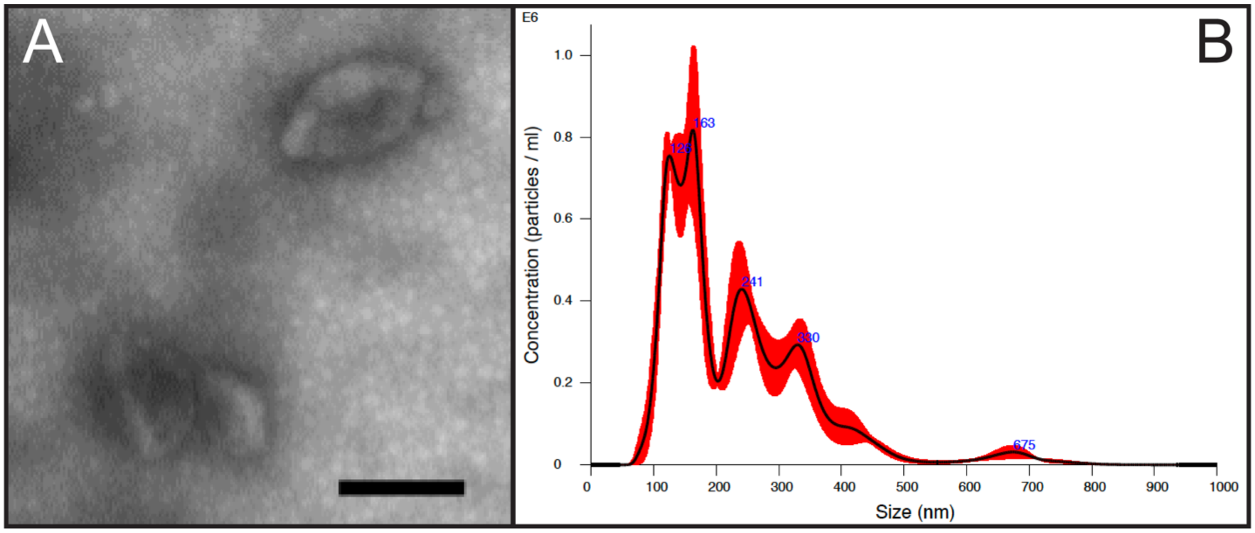


**Figure S3. Isolated EV Characterization. (A)** Isolated EVs were imaged by transmission electron microscopy (TEM) to confirm vesicle morphology. Representative TEM image shown for ultracentrifugation-isolated fibroblast EV sample. Scale bar = 200 nm. **(B)** Isolated EVs were analyzed with nanoparticle tracking analysis (NTA) to determine approximate particle size and concentration. Representative NTA data shown for ultracentrifugation-isolated fibroblast EV sample.
